# Mapping the lineage relationship between CXCR5^+^ and CXCR5^-^ CD4^+^ T cells in HIV infected human lymph nodes

**DOI:** 10.1101/537050

**Authors:** Daniel Del Alcazar, Yifeng Wang, Chenfeng He, Ben S. Wendel, Perla M. Del Río-Estrada, Jerome Lin, Yuria Ablanedo-Terrazas, Michael J Malone, Stefany M. Hernandez, Ian Frank, Ali Naji, Gustavo Reyes-Terán, Ning Jiang, Laura F. Su

**Affiliations:** Department of Medicine, Division of Rheumatology, Philadelphia VA Medical Center, University of Pennsylvania Perelman School of Medicine, Philadelphia, PA 19104, USA; Institute for Immunology, University of Pennsylvania Perelman School of Medicine, PA 19104, USA; Laboratory of Systems Immunology, Department of Biomedical Engineering, Cockrell School of Engineering, University of Texas at Austin, Austin, TX 78712, USA; McKetta Department of Chemical Engineering, Cockrell School of Engineering, University of Texas at Austin, Austin, TX 78712, USA; Departamento de Investigación en Enfermedades Infecciosas, Instituto Nacional de Enfermedades Respiratorias, Ciudad de México, México; Institute for Cellular and Molecular Biology, College of Natural Sciences, University of Texas at Austin, Austin, TX 78712, USA; Department of Oncology, Dell Medical School, University of Texas at Austin, Austin, TX 78712, USA; Institute for Biomedical Informatics, University of Pennsylvania, Philadelphia, PA 19104, USA; Department of surgery, University of Pennsylvania Perelman School of Medicine, Philadelphia, PA 19104, USA; Department of Medicine, Division of Infectious Disease, University of Pennsylvania Perelman School of Medicine, Philadelphia, PA 19104, USA

## Abstract

CXCR5 is a key surface marker expressed on follicular helper T (T_FH_) cells. We report here B cell help functionality in a population of CD4^+^ T cells isolated from primary human lymph nodes (LN) that lacked CXCR5 expression. This CXCR5^-^ subset is distinguished from other CXCR5^-^ CD4^+^ T cells by high PD-1 expression. Accumulation of CXCR5^-^PD-1^+^ T cells correlated with peripheral CD4^+^ T cell depletion and an increase in T-bet^+^ B cells in the LN, highlighting these atypical CD4^+^ T cells as a key component of lymphoid dysregulation during chronic HIV infection. By interrogating the phenotypic heterogeneity, functional capacity, TCR repertoire, transcriptional profile, and epigenetic state of CXCR5^-^PD-1^+^ T cells, we showed that CXCR5^-^PD-1^+^ T cells are related to CXCR5^+^PD-1^+^ T cells and provided evidence for the down regulation of CXCR5 following cell division as one mechanism for the absence of CXCR5 expression. Notably, CXCR5^-^PD-1^+^ T cells exhibited a migratory transcriptional program and contributed to circulating CXCR5^-^PD-1^+^ T cells with B cell help functionality in the peripheral blood. Thus, these data link LN pathology to circulating T cells and expand the current understanding on T cell diversity in the regulation of B cell responses during chronic inflammation.

- High dimensional profiling of activated CD4^+^ T cells in HIV infected lymph nodes revealed an accumulation of a CXCR5 negative subset.
- CXCR5^-^PD-1^+^CD4^+^ T cells exhibited T_FH_-like protein expression and function.
- CXCR5^-^PD-1^+^CD4^+^ T cells are related to T_FH_ cells by clonal lineage and epigenetic similarity.
- CXCR5^-^PD-1^+^CD4^+^ T cells upregulate a migratory gene program and contribute to circulating T cells with B cell help functionality

## Introduction

T cell activation is a hallmark of chronic HIV infection (Hunt et al., 2016; Sereti and Altfeld, 2016). T cells from HIV^+^ patients express increased levels of activation markers, CD38 and HLA-DR, which predict more rapid progression to AIDS in advanced HIV infection (Balagopal et al., 2015; Giorgi et al., 1993; Karim et al., 2013; Langford et al., 2007). Even with effective anti-retroviral therapy, T cell activation remains elevated in HIV infected individuals, likely as a result of viral persistence (Hunt et al., 2016; Lorenzo-Redondo et al., 2016). Lymphoid tissues are a major reservoir of HIV infection (Hufert et al., 1997; Kohler et al., 2016). Viral infection leads to disrupted lymphoid architectures and altered cellular differentiation (Hong et al., 2016). In particular, studies of human primary lymph nodes (LN) from untreated HIV patients have revealed an expansion of follicular helper T cells (T_FH_) (Lindqvist et al., 2012; Matthieu et al., 2013), which are classically identified by the expression of CXCR5, a chemokine receptor that enables proper follicular localization in the LN (Crotty, 2014; Haynes et al., 2007). T_FH_ cells are necessary for the affinity maturation process of B cells to generate high-affinity and broadly neutralizing antibodies (Havenar-Daughton et al., 2017). However, in spite of an increase in the abundance of T_FH_ cells in the LN, protective antibody responses to vaccines are generally diminished in the setting of HIV infection (Crum-Cianflone et al., 2011; de Armas et al., 2017). T_FH_ cells from HIV^+^ patients acquire a skewed functional phenotype and limited TCR diversity under persistent antigen stimulation (Wendel et al., 2018). Functional assays performed *in vitro* also showed provision of less effective B cell help by T_FH_ cells from HIV infected LNs (Cubas et al., 2013).

Due to the importance of T_FH_ cells in generating protective antibody responses, there have been substantial efforts to understand and manipulate T_FH_ cells for better vaccine efficacy. By comparison, much less is known about other cell types in inflamed LNs. Because HIV-driven immune hyperactivation extends to T cells in lymphoid tissues (Biancotto et al., 2007), we hypothesized that a more comprehensive understanding the complexity of activated T cells in the lymphoid compartment could provide insights into dysregulated T: B cell interactions for the improvement of efficacious protective antibody responses. In this study, we started by applying high-dimensional mass cytometry (CyTOF) to examine activated T cells in primary human LNs from HIV infected individuals, with the goal of discovering T cell populations that contribute to abnormal T cell responses in the lymphoid environment during chronic viral infection.

Here, we describe the identification of an activated CXCR5^-^ CD4^+^ T cell population that correlated with markers of HIV disease severity. These CXCR5^-^ CD4^+^ T cells are characterized by high PD-1 expression and are notable for their resemblance to T_FH_ cells by overlapping TCR repertoire sequences and similarities in transcriptional and epigenetic profiles. Importantly, the CXCR5^-^PD-1^+^ phenotype contains T cells that are capable of driving plasma cell differentiation and antibody production in response to stimulation by HIV antigens. To address how CXCR5^-^PD-1^+^ T cells differ from CXCR5^+^PD-1^+^ T cells, we compared their gene expression and identified a strong migratory signature involving upregulation of genes for LN egress and lymphocyte chemotaxis in CXCR5^-^PD-1^+^ T cells. Analyses of CD4^+^ T cells in peripheral blood mononuclear cells (PBMC) showed the accumulation of a related CXCR5^-^PD-1^+^ T cell population with B cell help functionality that shared overlapping TCR repertoire sequences with CXCR5^-^PD-1^+^ T cells from the LN. Collectively, these data provide an overview of the activated CD4^+^ T cells in HIV infected LNs and generate key insights into the nature of CD4^+^ T cell help for B cells in the setting of chronic inflammation.

## Results

### High dimensional analysis revealed an accumulation of activated CXCR5^-^CD4^+^ T cells in HIV infected lymph nodes

T cell activation is a hallmark of chronic HIV infection that predicts increased mortality in severe HIV infection (Giorgi et al., 1999). To better understand the heterogeneity of activated T cells in the LN, we performed mass cytometry with a 36-marker panel using lymph nodes from eight virally active HIV^+^ patients. Cryopreserved LN cells were stimulated with phorbol-12-myristate-13-acetate (PMA) and ionomycin in the presence of brefeldin A and monensin for 5 hours, stained with metal-conjugated antibodies, and analyzed on the mass cytometer, CyTOF 2. Data normalization was performed using a bead-based standards to minimize variations due to batch and machine performance (Finck et al., 2013). We defined activated T cells by CD38 and HLA-DR expression and compared CD38 and HLA-DR double positive T cells to quiescent CD3^+^ T cells that were negative for CD38 and HLA-DR staining (Fig. S1A). Activated or quiescent CD3^+^ T cells were further divided into αβ (CD4^+^ or CD8^+^) or γδ T cells by TCR staining. We showed that the majority of CD3^+^ cells expressed αβ TCRs (Fig. 1A). The CD4^+^ subset dominated the CD38^-^HLA-DR^-^ subset, but there was a notable shift toward CD8^+^ T cells among CD38^+^HLA-DR^+^ T cells (Fig. 1A and S1A). Intriguingly, the frequency of activated CD4^+^ T cells appeared to be better preserved in patients with more severe HIV infection as measured by an inverse correlation between CD38^+^HLA-DR^+^CD4^+^ T cell frequency and CD4:CD8 ratio in the blood (Fig. 1B). These data showed an accumulation of activated T cells in the LNs and suggested the possibility that CD4^+^ T cell depletion may be masked by the expansion of certain activated abnormal CD4^+^ T cells in LNs from patients with more severe HIV infection.

**Figure 1:**
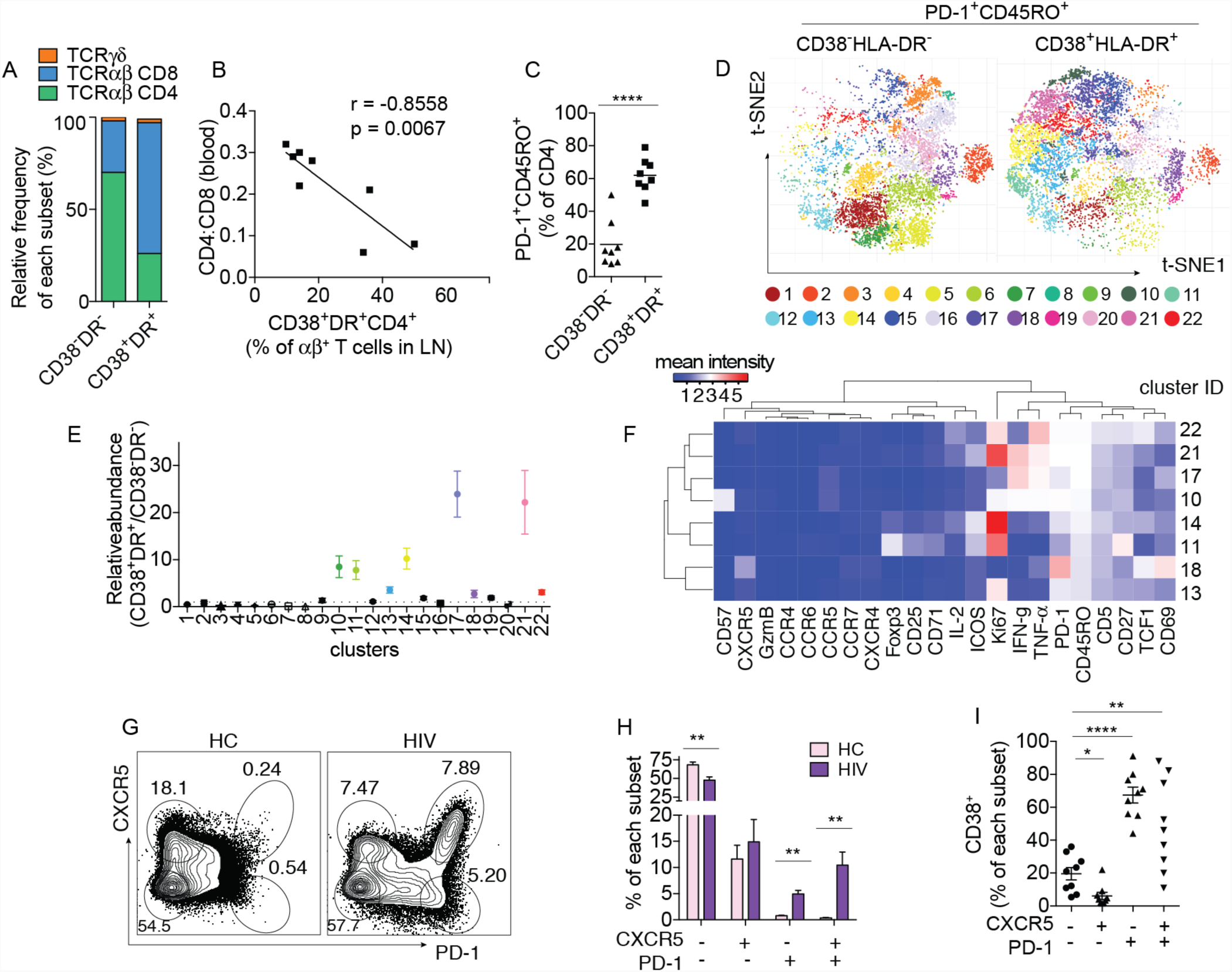
High dimensional analyses of activated T cells in HIV infected LNs by mass cytometry. A-F: CyTOF analyses of 8 HIV infected LNs. (A) Bar-graph shows the relative contribution of major T cell subsets in HIV infected LNs by TCR and coreceptor expression. (B) Correlation between peripheral CD4:CD8 ratio and the frequency of CD38^+^HLA-DR^+^CD4^+^ T cells in the LNs. (C) Quantification of PD-1^+^ memory T cells within active versus quiescent T cell subsets. (D) PD-1^+^CD45RO^+^CD4^+^ T cells gated by CD38^-^HLA-DR^-^ or CD38^+^HLA-DR^+^ phenotype were pooled and analyzed using “cytofkit” package in R. Data visualization is displayed on a two-dimensional t-SNE map. Twenty-two distinct phenotypic clusters were identified from PD-1^+^ memory CD4^+^ T cells using PhenoGraph. Also see Fig. S1. (E) Plot shows the frequency of each cluster in CD38^+^HLA-DR^+^ versus CD38^-^HLA-DR^-^CD4^+^ T cells (CD38^+^HLA-DR^+^ /CD38^-^HLA-DR^-^). A ratio of 2.5 is used as a cutoff to select clusters enriched in activated CD4^+^ T cells (colored). (F) Heatmap shows the average staining for markers that yielded positive signals (2 standard deviation above CD8 background) in at least one of the 8 populations. Markers used to select input cells were excluded. Data combine 8 HIV infected LNs and include both CD38^+^HLA-DR^+^ and CD38^-^HLA-DR^-^ CD4^+^ T cells. G-I: Flow cytometry analyses of 7 HC and 9 HIV LNs. (G-H) Example plots showing CXCR5 and PD-1 staining of CD4^+^ T cell. Bar-graph quantifies the frequency of each phenotypic subset. (I) The frequency of CD38^+^ T cells within each CD4^+^ T cell subset. For (B), association was measured by Pearson correlation and the best-fitting line was calculated using least squares fit regression. For (C), paired t-test was used. For (H), multiple t-tests comparisons were performed and corrected using Holm-Sidak method. For (I), one-way ANOVA was performed and corrected using Dunnett’s multiple comparisons test.

To identify CD4^+^ T cells that were over-represented in the activated subset, we focused on cells expressing inhibitory receptor PD-1, which was detected in over 60% of CD38^+^HLA-DR^+^CD4^+^ T cells (Fig. 1C). PD-1^+^CD45RO^+^CD4^+^ T cells from the CD38^+^HLA-DR^+^ and CD38^-^HLA-DR^-^ subsets of eight HIV infected LNs were combined and partitioned into clusters of phenotypically similar cells using PhenoGraph (Levine et al., 2015). Data visualization was performed using T-Distributed Stochastic Neighbor Embedding (t-SNE), which clearly showed distinct patterns between CD38^+^HLA-DR^+^ and CD38^-^HLA-DR^-^ T cells (Fig. 1D). To determine which clusters were more abundant in activated CD4^+^ T cells, we divided the frequency of each cluster in CD38^+^HLA-DR^+^ T cells by its frequency in the CD38^-^HLA-DR^-^ subset and used over 2.5-fold enrichment as a criteria for selecting clusters that were preferentially enriched in activated T cells for further analyses (Fig. 1E). In total, we identified 8 clusters of interest, which contributed to over 60% of activated CD4^+^ T cells on average (Fig. 1E, Fig. S1B). Using a heatmap to display the staining intensity for positive staining markers (2 standard deviation above the CD8 background) and excluding those that were used to select the input cells, we found only one out of the 8 activation-enriched clusters stained brightly for CXCR5 (Fig. 1F). These data suggest that the expansion of activated CD4^+^ T cells in HIV infected LNs is not limited to T_FH_ cells and included other types of CD4^+^ T cells. We performed fluorescent cytometry on a separate set of LN cells to confirm the findings above and found an increase in both CXCR5 positive and negative subsets of PD-1^+^ cells in HIV infected LNs compared to cells from healthy controls (HCs) (Fig. 1G-H). Both PD-1^+^ phenotypic subsets, irrespective of the level of CXCR5 expression, were enriched for activated T cells by CD38 staining (Fig. 1I). Taken together, our data confirmed the expansion of T_FH_ cells in HIV infected LNs and revealed the accumulation of another activated PD-1^+^ population that lacked CXCR5 expression during chronic HIV infection.

### CXCR5^-^PD-1^+^ T cells express exhaustion-related surface markers but retain cytokine producing potential

We were interested in how the expansion of a CXCR5^-^PD-1^+^ T cell population relates to clinical parameters of HIV infection. Using a previously acquired dataset from 25 HIV infected LNs (Wendel et al., 2018), we showed that the frequency of CXCR5^-^PD-1^+^ T cells in the LNs inversely correlated with CD4:CD8 ratio and CD4^+^ T cell count in the peripheral blood, but not with the viral load (Fig. 2A and Fig. S2A-B). We hypothesized that PD-1 expression on these cells reflected T cell exhaustion, possibly driven by a combination of the depleted CD4^+^ T cell niche and chronic antigen stimulation by HIV infection. To test this, we stained HIV infected LN cells for other exhaustion related surface molecules, LAG3, TIGIT, 2B4, TIM3, and CD39 (Crawford et al., 2014; Simoni et al., 2018). We defined non-exhausted memory cells based on the absence of PD-1 staining and compared the expression of LAG3, TIGIT, 2B4, TIM3, and CD39 as a percentage of PD-1^+^ or PD-1^-^ CD45RO^+^ T cells. CXCR5 expressing CD4^+^ cells were initially excluded to focus on the non-T_FH_ population. Compared to the PD-1^-^ subset, we observed a significant increase in LAG3 and TIGIT expression on PD-1^+^ cells (Fig. 2B and S3A). We next focused on the differences between CXCR5^-^ and CXCR5^+^ cells within PD-1^+^ population. This revealed a trend for higher TIGIT and 2B4 expression on the CXCR5^+^ T cells and significant increase in CD39 staining on the CXCR5^-^ subset (Fig. 2C and S3B).

**Figure 2:**
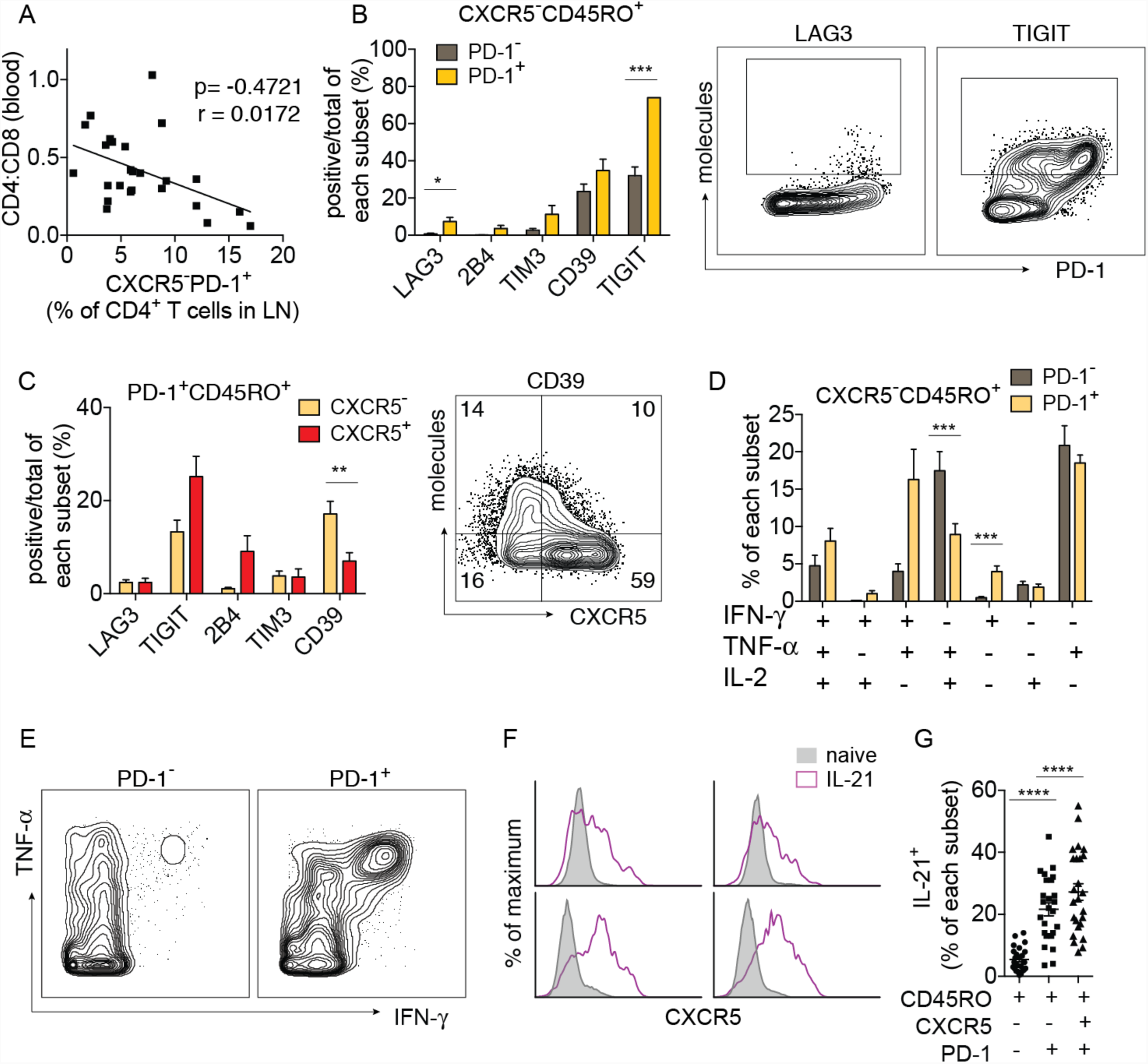
CXCR5^-^PD-1^+^ T cells express exhaustion-related surface markers but retain cytokine producing potential. (A) Correlation between peripheral CD4:CD8 ratio and the frequency of CXCR5^-^PD-1^+^subset of CD4^+^ T cells in the LNs (n = 25 HIV^+^ donors). Also see Fig. S2A-B. (B) Bar-graph showing the relative expression of the indicated exhaustion-related markers as a percentage of PD-1^-^ or PD-1^+^ subset of CXCR5^-^ memory CD4^+^ T cells (n = 8 HIV LNs). Plots show representative staining of differentially expressed markers. (C) A comparison of exhaustion-related markers in CXCR5^-^ versus CXCR5^+^ memory subsets (n = 8 HIV LNs). Plot shows representative staining for CD39. Also see Fig. S3. (D) The frequency of PD-1^-^ or PD-1^+^ cells that positively stained for the indicated cytokines after 5 hours of PMA and ionomycin stimulation as determined by CyTOF. CXCR5^+^ cells were excluded (n = 8 HIV LNs). Also see Fig. S4A. (E) Plots showing representative staining for IFN-γ and TNF-α in PD-1^-^ and PD-1^+^ subset of CXCR5^-^ memory CD4^+^ T cells. (F) LN cells from 4 HIV^+^ donors were stimulated with PMA and ionomycin and assayed by flow cytometry for IL-21 expression. Histograms compare the fluorescent intensity of CXCR5 staining in IL-21^+^ T cells versus naïve T cells. Each histogram represents data from one donor. Also see Fig. S4B. (G) Frequency of IL-21^+^ T cells in each indicated subset following PMA and ionomycin stimulation as measured by CyTOF (n = 25 HIV LNs). Also see Fig. S4C. For (A), association was measured by Pearson correlation and the best-fitting line was calculated using least squares fit regression. For (B), (C), and (D), multiple t-tests comparisons were performed and multiple comparison was corrected using Holm-Sidak method. For (G), one-way ANOVA was performed and corrected using Dunnett’s multiple comparisons test.

It was unclear if upregulation of exhaustion-related markers was indicative of functional exhaustion. To examine the functional capacity for cytokine production, Boolean combination gates defining all possible combinations of TNF-α, IFN-γ, and IL-2 staining were applied to PD-1^+^ and PD-1^-^ memory T cells stimulated with PMA and ionomycin (Fig. S4A). We found that PD-1^+^ T cells have the potential to produce significantly more IFN-γ upon T cell activation (Fig. 2D-E). Compared to PD-1^-^ T cells, fewer PD-1^+^ T cells were IL-2^+^TNF-α^+^ double producers but both subsets contained similar frequencies of polyfunctional T cells that produce all three cytokines (Fig. 2D-E). Because IL-21 is a critical cytokine in the lymphoid environment, we next asked if PD-1^+^ T cells deficient in CXCR5 expression were capable of making IL-21. We first examined IL-21 staining on PMA and ionomycin stimulated T cells by flow cytometry and found no clear restriction of IL-21 expression to cells that stained more strongly for CXCR5 (Fig. 2F and S4B). We confirmed this observation by reanalyzing the 25-sample CyTOF dataset, which we had previously used for a focused T_FH_ cell analysis (Wendel et al., 2018). In comparison with memory cells with a CXCR5^-^PD-1^-^ phenotype, the frequency of IL-21 producing CXCR5^-^PD-1^+^ T cells was significantly higher and approached that of CXCR5^+^PD-1^+^ T cells (Fig. 2G and S4C). Thus, PD-1 expression on CXCR5^-^ T cells in HIV infected LNs marked a population of functionally active T cells with a unique cytokine profile.

### CXCR5^-^PD-1^+^ T cells expressed T_FH_ cell-related proteins and exhibit similar activity in vitro

The ability to produce IL-21 under stimulation suggested the possibility that this CXCR5 negative population may be more similar to T_FH_ cells than exhausted T cells or other types of conventional CD4^+^ T cells. To examine additional T_FH_ cell-related features in CXCR5^-^ T cells, we stained LN cells for BCL6, Maf, and CD84 expression. BCL6 and Maf are critical transcriptional regulators of T_FH_ cell specification and function (Bauquet et al., 2009; Yu et al., 2009). CD84 is a SLAM family receptor that transduces adhesion signals to drive germinal center differentiation (Cannons et al., 2010). We compared staining for BCL6, Maf, and CD84 in naïve, CXCR5^-^PD-1^-^, CXCR5^-^PD-1^+^, and CXCR5^+^PD-1^+^ T cells (Fig. S5). While BCL6 and Maf expression remained highest in CXCR5^+^PD-1^+^ T cells, all three proteins were significantly increased in CXCR5^-^PD-1^+^ compared to naïve T cells (Fig. 3A-B). To determine if the partial resemblance between CXCR5^-^PD-1^+^ and CXCR5^+^PD-1^+^ T cells extended to a similar functional program, we performed T:B coculture assays using sorted CXCR5^+^PD-1^+^ or CXCR5^-^PD-1^+^ T cells (Fig. S6A). We added ICOS to the sort strategy because ICOS and PD-1 were co-expressed and ICOS expression further enriched for cells with an activated phenotype (Fig. S6A-B). Including an ICOS gate also helped to better define CXCR5^+^ T cells of a classic T_FH_ phenotype. T cells were cultured with memory B cells at 1:1 ratio and stimulated with Staphylococcus aureus Enterotoxin B (SEB). CXCR5^-^PD-1^+^ T cells clearly support B cells in this experimental setting and induced similar levels of plasma cell differentiation and antibody production compared to CXCR5^+^PD-1^+^ T cells (Fig. 3C-D). Next, we tested the *in vivo* relevance of CXCR5^-^PD-1^+^ T cells by measuring correlative changes in B cell differentiation in primary LN samples. Because CXCR5^-^PD-1^+^ T cells secrete high levels of IFN-γ, which has been shown to induce T-bet expression in B cells (Naradikian et al., 2016), we focused our analyses on T-bet^+^ B cell subset. We identified T-bet^+^ B cells by T-bet and CXCR3 co-expression (Fig. 3E). Compared to cells from HCs, T-bet^+^ B cells were significantly increased in HIV infected LNs and their abundance correlated with the frequency of IFN-γ producing CXCR5^-^PD-1^+^ T cells (Fig. 3F-G). Collectively, these data suggest a role for CXCR5^-^PD-1^+^ T cells in the regulation of B cell responses during chronic HIV infection.

**Figure 3:**
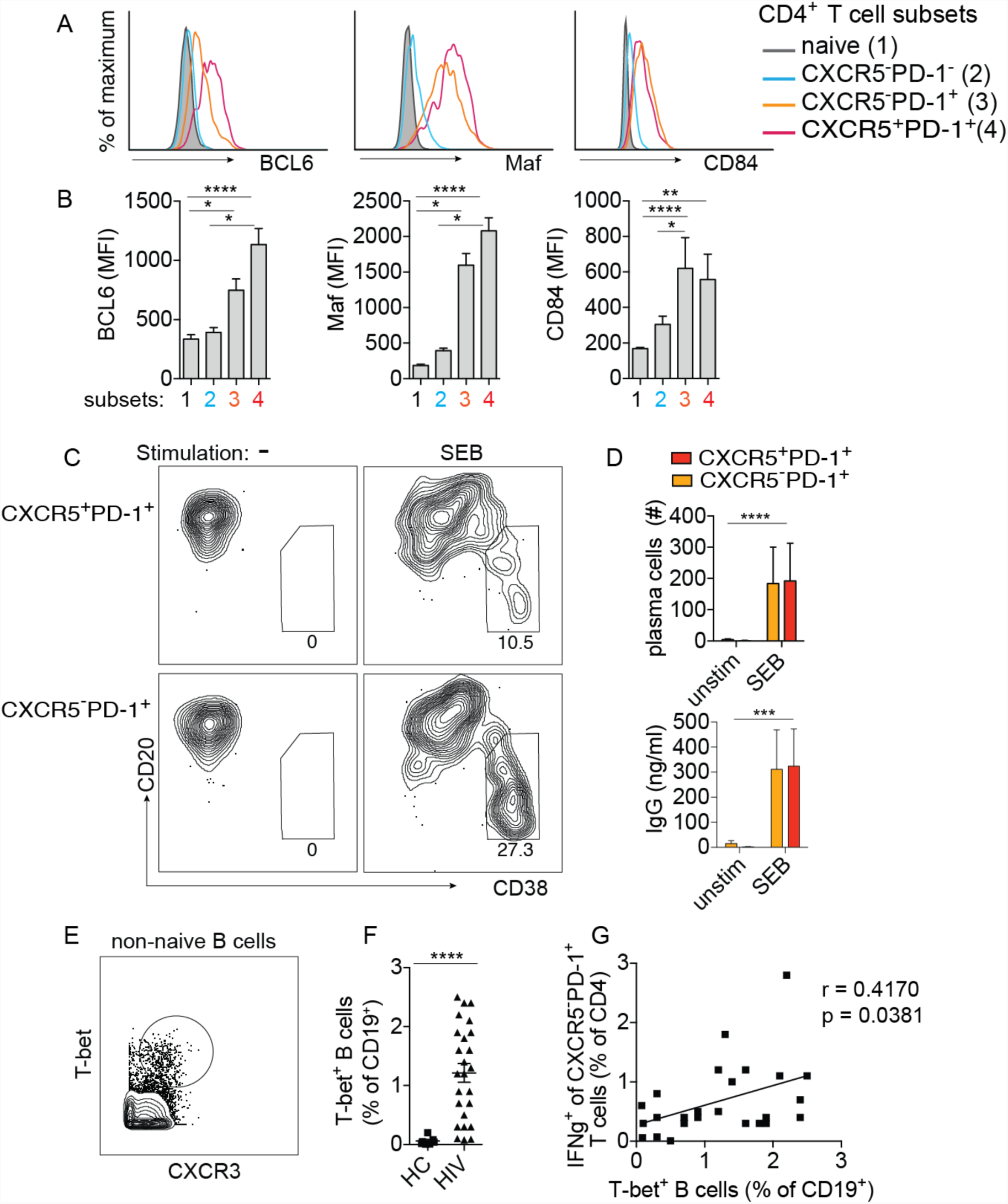
CXCR5^-^PD-1^+^ T cells expressed T_FH_ cell-related proteins and function. (A-B) Histograms showing fluorescent intensity of BCL6, Maf, and CD84 staining in naïve, CXCR5^-^PD-1^-^, CXCR5^-^PD-1^+^, and CXCR5^+^PD-1^+^ T cells (A). Data from 8 HIV LNs are summarized in (B). Also see Fig. S5. (C) Purified CXCR5^-^PD-1^+^ and CXCR5^+^PD-1^+^ T cells were cultured with memory B cells in the presence or absence of SEB for 7 days. Plots show representative staining for plasma cells. Also see Fig. S6. (D) Bar-graph quantifies plasma cell (top, n = 11 HIV LNs) and IgG (bottom, n = 9 HIV LNs) at the end of coculture. (E) Example Plot showing gating for T-bet^+^ B cells. Naïve IgD^+^CD27^-^ B cells were excluded. (F) Quantification of T-bet^+^ B cells in 7 HC and 25 HIV infected LNs. (G) Correlation between IFN*–*γ expression in CXCR5^-^PD-1^+^ T cells and T-bet^+^ B cell frequency in HIV infected LN samples. For (B), one-way ANOVA was performed and corrected using Dunnett’s multiple comparisons test. For (D), two-way ANOVA was performed and corrected using Sidak’s multiple comparisons test. For (F), Mann Whitney test was used. For (G), association was measured by Spearman correlation.

### CXCR5^-^PD-1^+^ T cells shared HIV-specificity and TCR sequences with CXCR5^+^PD-1^+^ T cells

Given the functional overlap between CXCR5^-^PD-1^+^ and CXCR5^+^PD-1^+^ T cells, we next asked if these phenotypically distinct populations shared a common clonal origin or if they developed independently from different T cell populations. We tracked the clonal relationship between CXCR5^-^PD-1^+^ T cells and CXCR5^+^PD-1^+^ T cells by taking advantage of T cell receptor (TCR) sequence as a unique T cell identifier. T cells expressing the identical TCR sequences are necessarily generated from the same precursors, and thus we can use TCR sequences to infer the relatedness between phenotypically distinct populations. We performed TCR repertoire sequencing on sorted naïve, CXCR5^-^PD-1^-^, CXCR5^-^PD-1^+^, and CXCR5^+^PD-1^+^ T cells (Fig. S6A). Common sequences shared between CXCR5^-^PD-1^+^ and other T cell subsets were shown as connecting lines on circos plots and quantified by the Bhattacharyya Coefficient as an index of sequence similarity (Bhattacharyya, 1943) (Fig. 4A). Analysis of the combined TCR sequencing data from 9 donors showed that the clonal overlap was highest between CXCR5^-^PD-1^+^ and CXCR5^+^PD-1^+^ T cells, which significantly exceeded the TCR sequence similarity between CXCR5^-^PD-1^+^ and naïve T cells (Fig. 4B).

**Figure 4:**
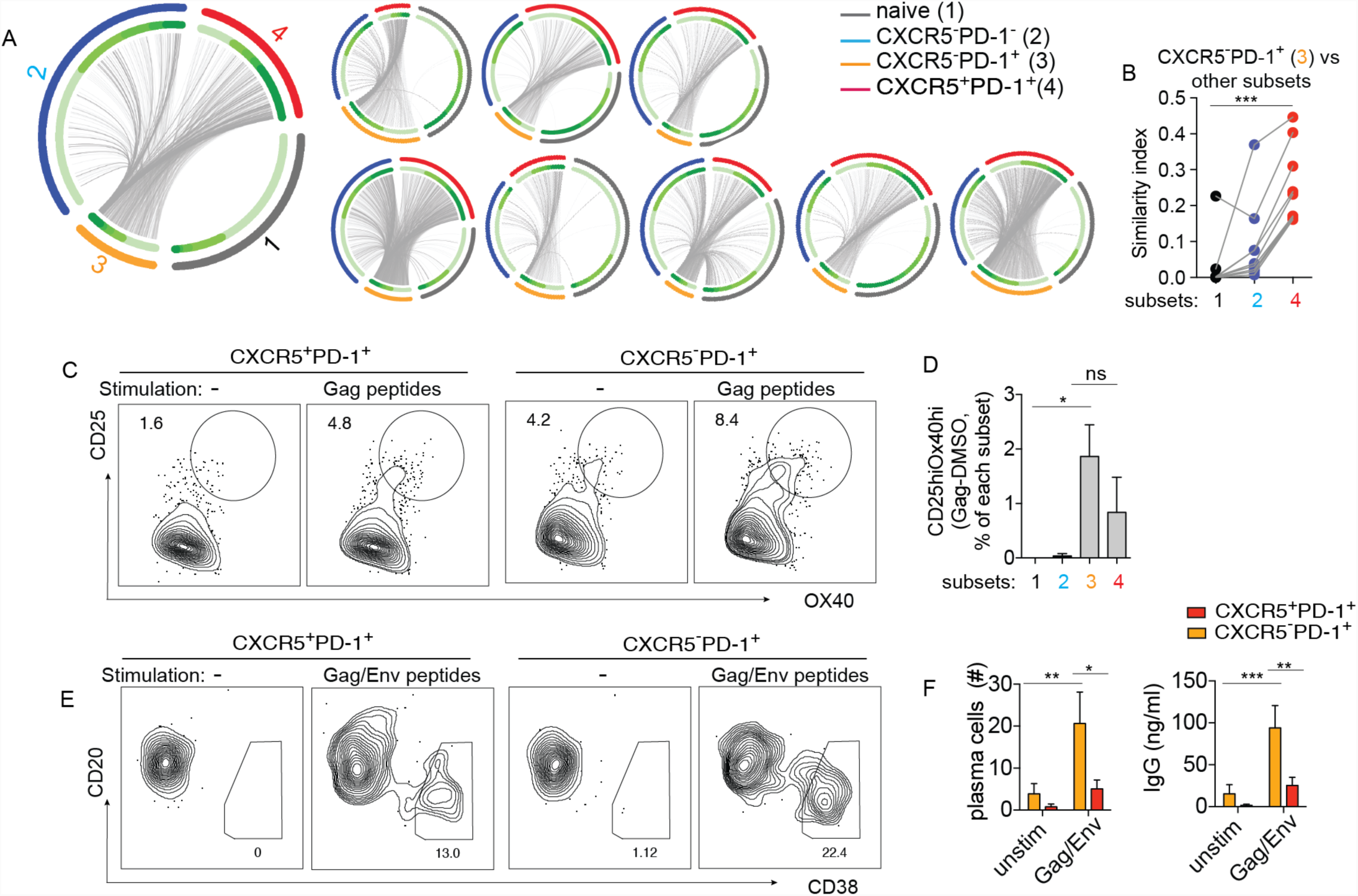
CXCR5^-^PD-1^+^ T cells shared HIV-specificity and TCR sequences with CXCR5^+^PD-1^+^ T cells. (A) Circos plots of TCR sequence overlap among different populations. Each thin slice of the arc represents a unique TCR sequence, ordered by the clone size (darker green for larger clones, inner circle). Outer circle indicate TCR nucleotide sequences found in naïve (gray), CXCR5^-^PD-1^-^ (blue), CXCR5^-^PD-1^+^ (orange), and CXCR5^+^PD-1^+^ (red) populations. Each plot represents data from one individual. (B) Bhattacharyya coefficient measurement for the TCR repertoire similarity between CXCR5^-^PD-1^+^ with other populations within LN. Grey lines connect samples from the same patient. Similarity index with naïve cells is used as the baseline to identify more related repertoires (n = 9 HIV LNs). (C) Gag-specific T cells within CXCR5^-^ and CXCR5^+^ subsets were identified by OX40 and CD25 expression in LN cells stimulated with pooled Gag peptides for 18 hours. Plots show representative data from one HIV LN. (D) Quantification of CD25hiOX40hi T cell frequency. Background level of CD25hiOX40hi T cells in vehicle-treated wells was subtracted from Gag-stimulated wells (n = 7 HIV LNs). (E) Sorted CXCR5^-^PD-1^+^ and CXCR5^+^PD-1^+^ T cells from HIV infected LNs were cultured with memory B cells in the presence or absence of pooled Env and Gag peptides for 7 days. Representative data from one individual is shown. (F) Bar-graph quantifies plasma cell (left, n = 11) and IgG (right, n = 9) at the end of coculture. Cocultures were performed in duplicates whenever possible with the available numbers of cells. For (D), one-way ANOVA was performed and corrected using Dunnett’s multiple comparisons test. For (F), two-way ANOVA was performed and corrected using Sidak’s multiple comparisons test.

Because T_FH_ cells are enriched for HIV-reactive T cells in HIV infected LNs (Lindqvist et al., 2012; Matthieu et al., 2013; Wendel et al., 2018), we tested if CXCR5^-^PD-1^+^ T cells are also enriched for T cells that recognize HIV-derived antigens. To do this, we stimulated total LN cells with Gag peptides for 18 hours and identified peptide-specific T cells within naïve, CXCR5^-^PD-1^-^, CXCR5^-^PD-1^+^, or CXCR5^+^PD-1^+^ T cells by CD25 and OX40 upregulation (Dan et al., 2016). The frequency of CD25 and OX40 double positive T cells in dimethyl sulfoxide (DMSO) vehicle treated backgrounds were subtracted from Gag peptide stimulated cultures to calculate the frequency of Gag-reactive T cells. On average, 1.86% of CXCR5^-^PD-1^+^ subset expressed OX40 and CD25 in response to peptide stimulation (range: 0-4.2%), whereas negligible numbers of CD25^+^OX40^+^ cells were detected in the naïve and CXCR5^-^PD-1^-^ subsets (Fig. 4C-D). To determine if HIV-reactive CXCR5^-^ PD-1^+^ T cells were functionally active, we stimulated cocultures of memory B cells and CXCR5^-^PD-1^+^ or CXCR5^+^PD-1^+^ T cells using overlapping pools of Gag and Env peptides instead of SEB. Induction of plasma cell differentiation and IgG production was detected in B cells cultured with peptide-stimulated CXCR5^-^PD-1^+^ T cells. Notably, the level of B cell response was higher in cultures containing CXCR5^-^PD-1^+^ T cells compared to CXCR5^+^PD-1^+^ T cells, consistent with the inclusion of a functional HIV-reactive population in the CXCR5 negative subset (Fig. 4E-F). Collectively, these data demonstrate the presence of a CXCR5^-^ CD4^+^ T cell population that is specific to HIV antigens and has the functional potential to impact B cell responses. The TCR repertoire sequencing data further suggest that this CXCR5^-^ subset is clonally related to CXCR5^+^PD-1^+^ T cells.

### Dynamic regulation of CXCR5 contributes to the accumulation of CXCR5^-^PD-1^+^ T cells in inflamed lymph nodes

To further define the relationship between CXCR5^-^PD-1^+^ and CXCR5^+^PD-1^+^ T cells, we used diffusion map to construct probable differentiation trajectories for the eight activation-enriched clusters identified by Phenograph/t-SNE in the CyTOF dataset (Fig. 1F). Diffusion map is a nonlinear dimension reduction tool that performs peudotemporal ordering of single cell data using the probability of transition between cells on a random walk (Haghverdi et al., 2015). It ordered the CXCR5^+^ cluster (cluster 18) at the beginning of the phenotypic progression and predicted a decrease in CXCR5 expression in cells progressing toward a more differentiated state (Fig. 5A). It accurately mapped low TCF1 expression to cells at the end of this peudotemperal progression, which has been experimentally shown to decrease with cell division (Nish et al., 2017) (Fig. 5A). Consistent with this differentiation trajectory, we manually gated for CXCR5 expression by TCF1 staining and demonstrated a clear partitioning of CXCR5^+^ T cells to the TCF1hi subset and low levels of CXCR5 staining in cells with low TCF1 expression (Fig. 5B). These data suggest that an initially CXCR5 expressing T cell may down regulate CXCR5 as it undergoes activation-induced cell division. To test this hypothesis, we labeled CXCR5^+^PD-1^+^ T cells with Carboxyfluorescein succinimidyl ester (CFSE). Dilution of CFSE was used to identify T cells that have divided after 7 days of culture in SEB. Compared to unstimulated T cells, T cells in SEB treated wells generally expressed lower levels of CXCR5, with a further reduction in CXCR5 staining for CFSE_lo_ T cells that had undergone cell division (Fig. 5C-D). To determine the *in vivo* relevance of this observation, we used the assay for transposase-accessible chromatin with sequencing (ATAC-seq) to provide a glimpse into the differentiation steps of CXCR5^-^PD-1^+^ T cells. An open chromatin region is predictive of permissive transcriptional state, and CXCR5^-^PD-1^+^ T cells are expected to have an open chromatin around CXCR5 promoter region if these cells had previously expressed CXCR5. Alternatively, we expect to find a closed CXCR5 promoter region if CXCR5^-^PD-1^+^ T cells did not go through a CXCR5^+^ state or if changes in the chromatin structure were short-lived. We generated a chromatin accessibility map for naïve, CXCR5^-^PD-1^-^, CXCR5^-^PD-1^+^, and CXCR5^+^PD-1^+^ T cells. Principal component analyses (PCA) mapped CXCR5^-^PD-1^+^ close to CXCR5^+^PD-1^+^ T cells on the first two PCA components, which accounted for the majority of the variation in this dataset (Fig. 5E). Focusing on CXCR5, we observed an accessible open chromatin state around the CXCR5 promoter region in CXCR5^-^PD-1^+^ T cell (Fig. 5F). Collectively, these data provide evidence for down regulation of CXCR5 expression in mature dividing CXCR5^+^ T cells and suggests a contribution of this process to the accumulation of CXCR5^-^PD-1^+^ T cells in inflamed LNs.

**Figure 5:**
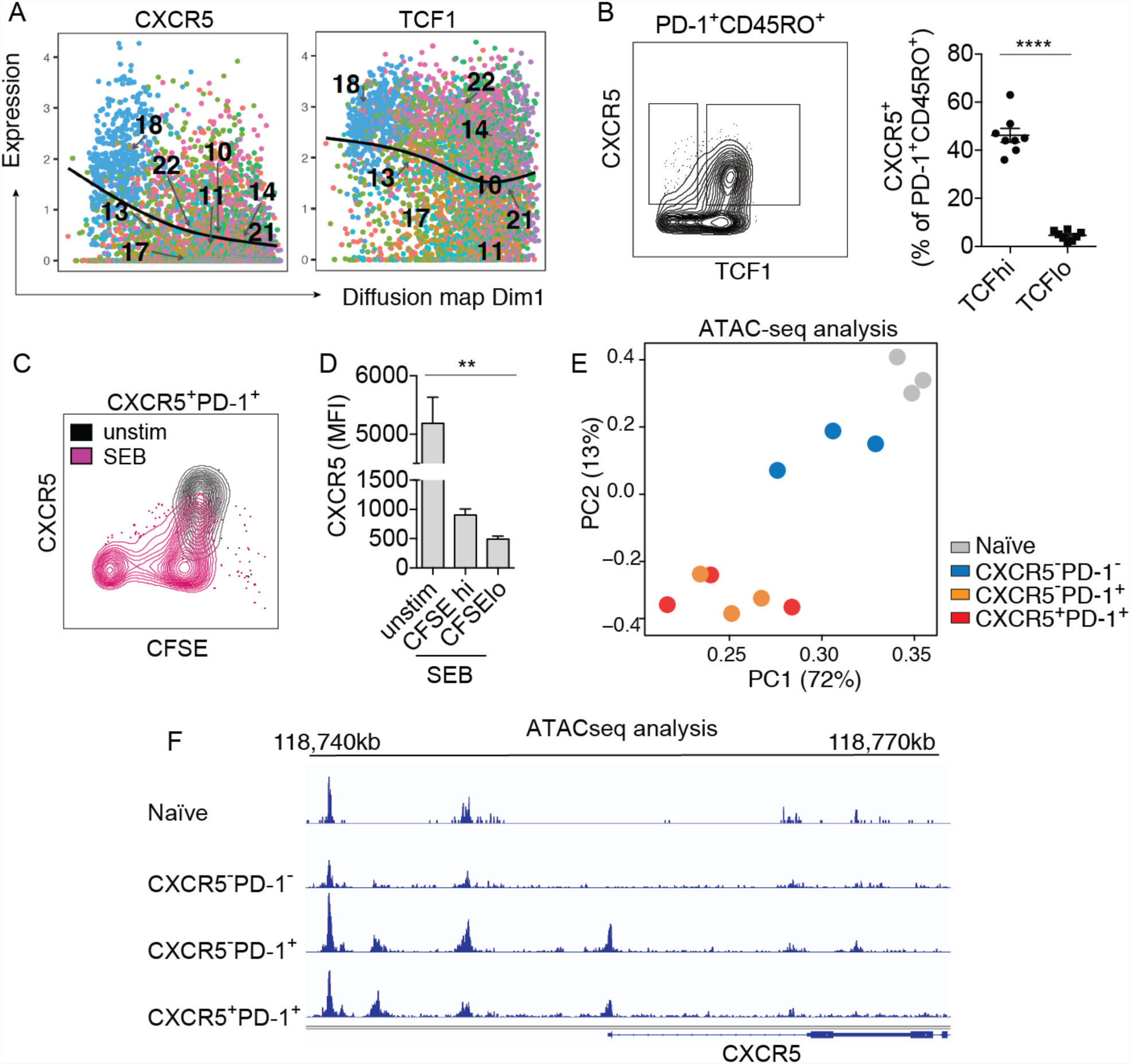
Dynamic regulation of CXCR5 contributed to the accumulation of CXCR5^-^PD-1^+^ T cells in inflamed LNs. (A) Hypothetical progression of the clusters that were significantly enriched (> 2.5 fold) in activated CD4^+^ T cells were ordered using diffusion map. The lines indicate the expression level of CXCR5 and TCF1 in pseudotime. (B) Example plot showing TCF1 and CXCR5 staining on PD-1^+^CD45RO^+^CD4^+^ T cells. Quantification of CXCR5^+^ T cells in memory subsets expressing high or low levels of TCF1 (n = 8 HIV LNs). (C) Sorted CXCR5^+^PD-1^+^ T cells were labeled with CFSE and stimulated with SEB or DMSO for 7 days in the presence of memory B cells. Plot shows representative CFSE and CXCR5 staining after coculture. (D) Bar-graph quantifies the fluorescence intensity of CXCR5 staining (n = 6). (E) Principal component analysis of ATAC-seq data from different CD4^+^ T cell subsets. Each symbol represent cells from one donor, cells of the same type are coded by the same color. (F) Representative example of chromatin accessibility at the CXCR5 upstream region for each CD4^+^ T cell subset. For (B), paired t-test was performed. For (D), Friedman test was performed and corrected using Dunn’s multiple comparisons test. Also see Fig. S6.

### CXCR5^-^PD-1^+^ T cells express a migratory gene program and contribute to CXCR5^-^PD-1^+^ T cells in the peripheral blood

To further elucidate the functional program of CXCR5^-^ T cells and understand how this population differs from CXCR5^+^ T cells, we performed transcriptomic profiling of Naïve, CXCR5^-^PD-1^-^, CXCR5^-^PD-1^+^, and CXCR5^+^PD-1^+^ T cells by RNA sequencing. Analyses of the RNA sequencing dataset showed marked similarity between CXCR5^-^PD-1^+^ and CXCR5^+^PD-1^+^ T cells. Similar to the PCA plots generated using the ATAC-seq data, CXCR5^-^PD-1^+^ T cells mapped closely to CXCR5^+^PD-1^+^ T cells in a distinct PCA space separated from naïve and CXCR5^-^PD-1^-^ T cells (Fig. 6A). To identify gene programs that are unique to CXCR5^-^PD-1^+^ T cells, we used gene set enrichment analysis (GSEA) to identify the GO biological pathways enriched in CXCR5^-^PD-1^+^ T cells. In parallel, we identified 221 genes differentially expressed between CXCR5^-^PD-1^+^ and CXCR5^+^PD-1^+^ T cells using DEseq2. We then used GOrilla to find GO biological pathways enriched by this gene set. The results from GSEA and differential gene expression analyses were combined to identify pathways common to both analyses methods. This focused the data on six CXCR5^-^PD-1^+^ T cell-upregulated pathways, four of which were involved in chemokine sensing and cell movement (Fig. 6B). Differentially expressed genes contributing to enriched GO terms were displayed in a heatmap, which included numerous membrane receptors involved in cell adhesion and migration (CCR2, CCR5, ITGB1, CXCR6, SELPLG, S1PR1) (Fig. 6C). Consistent with a T_H1_-skewed cytokine profile, T-bet and Runx3 expressions were increased in CXCR5^-^PD-1^+^ T cells (Djuretic et al., 2007; Szabo et al., 2000). Expectedly, the absence of CXCR5 protein expression correlated with low levels of CXCR5 mRNA expression. Other T_FH_ cell-associated transcripts, BCL6 and CXCL13, were also lower in CXCR5^-^PD-1^+^ T cells compared to CXCR5^+^PD-1^+^ T cells. Thus, CXCR5^-^PD-1^+^ T cells appear to be a transitional population that is losing gene signatures of T_FH_ cells while acquiring new functional characteristics.

**Figure 6:**
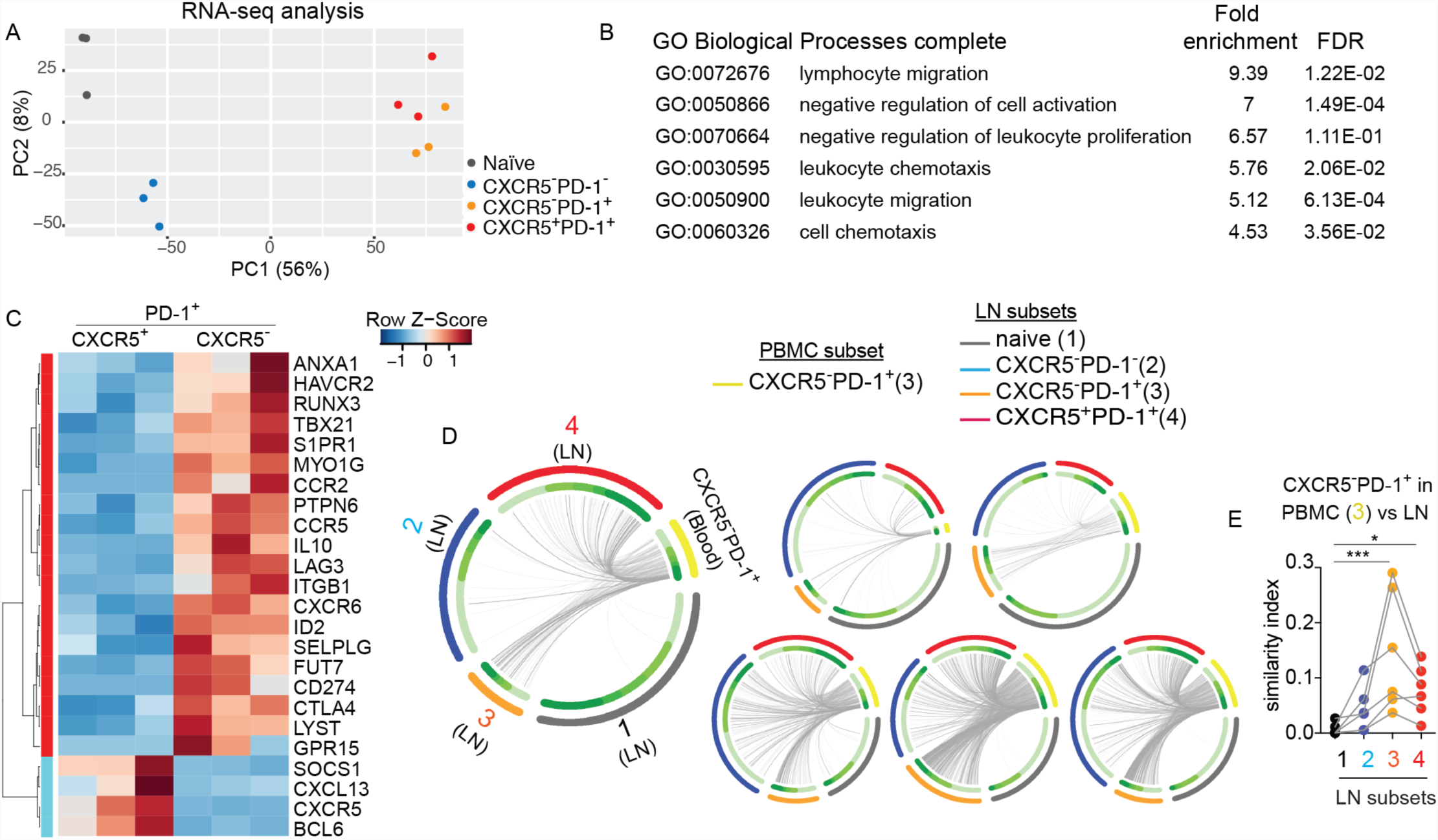
Migration-related gene signatures and PBMC-derived TCRs are enriched in CXCR5^-^PD-1^+^ T cells from HIV infected LNs. (A) Principal component analysis of RNAseq data from different CD4^+^ T cell subsets. Each symbol represent cells from one donor, cells of the same type are coded by the same color. (B) A list of gene ontologies based on significantly variant genes compared between CXCR5^-^PD-1^+^ / CXCR5^+^PD-1^+^ T cells. The list is a collection of hits called by both GSEA and GOrilla. (C) Heatmap for supervised clustering of 24 differentially expressed genes that contributed to the enriched GO terms in (B). The color scale represents the scaled normalized expression value as z-scores. (D) Circos plots of TCR sequence overlap among different populations. Each thin slice of the arc represents a unique TCR sequence. CXCR5^-^PD-1^+^ from PBMC is indicated in yellow. Each plot represents data from one individual. (E) Bhattacharyya coefficient measurement for the TCR repertoire similarity between CXCR5^-^PD-1^+^ in the PBMC and the indicated T cell subsets from the LN. Similarity index with naïve cells is used as the baseline to identify more related repertoires. For (E), Friedman test was performed and corrected using Dunn’s multiple comparisons test. Also see Fig. S6.

S1PR1 responds to high concentration of S1P in circulating fluid to guide lymphocytes out of the lymph nodes (Cyster and Schwab, 2012). To test if migration-related signature in CXCR5^-^PD-1^+^ T cells promoted their egress from the LN, we sorted CXCR5^-^PD-1^+^ T cells in the PBMC and compared TCR sequences from PBMC and LN cells from the same donors (Fig. S6D). We hypothesize that we can capture identical TCR sequences in distinct compartments if there are intermixing between cells in blood and LN. A comparison of the TCR repertoires showed the similarity index were indeed the highest between the TCR repertoires of blood and LN-derived CXCR5^-^PD-1^+^ T cells (Fig. 6D-E). In addition, TCRs expressed by circulating CXCR5^-^PD-1^+^ T cells also shared a significant overlap with the TCRs of CXCR5^+^PD-1^+^ T cells in the LN (Fig. 6D-E), suggesting a direct or indirect contribution from the CXCR5^+^PD-1^+^ lymphoid subset.

Next, we quantified the frequency of CXCR5^-^PD-1^+^ T cells in PBMC and compared it with different T cell subsets in the LN. We found relatedness by TCR repertoire sequencing predicted shared abundance across compartments. Where there were similarities in the TCR repertoires, the frequency of circulating CXCR5^-^PD-1^+^ T cells correlated with that of CXCR5^-^PD-1^+^ and CXCR5^+^PD-1^+^ T cells in the LN. In contrast, CXCR5^-^PD-1^+^ T cell frequency in the blood did not correlate with the abundance of lymphoid CXCR5^-^PD-1^-^ T cells, with which there was little TCR overlap (Fig. 6E and 7A). Similar to CXCR5^-^PD-1^+^ T cells in the LN, the abundance of CXCR5^-^PD-1^+^ T cells in the blood was predictive of more severe HIV infection as measured by a lower CD4^+^ T cell count and CD4:CD8 ratio (Fig. 7B and S2C-D). Thus, populations that share similar TCR repertoires exhibited linked frequencies, thereby enabling detection of CXCR5^-^PD-1^+^ T cells in the PBMC to inform changes in the representation of related populations in the LN.

**Figure 7:**
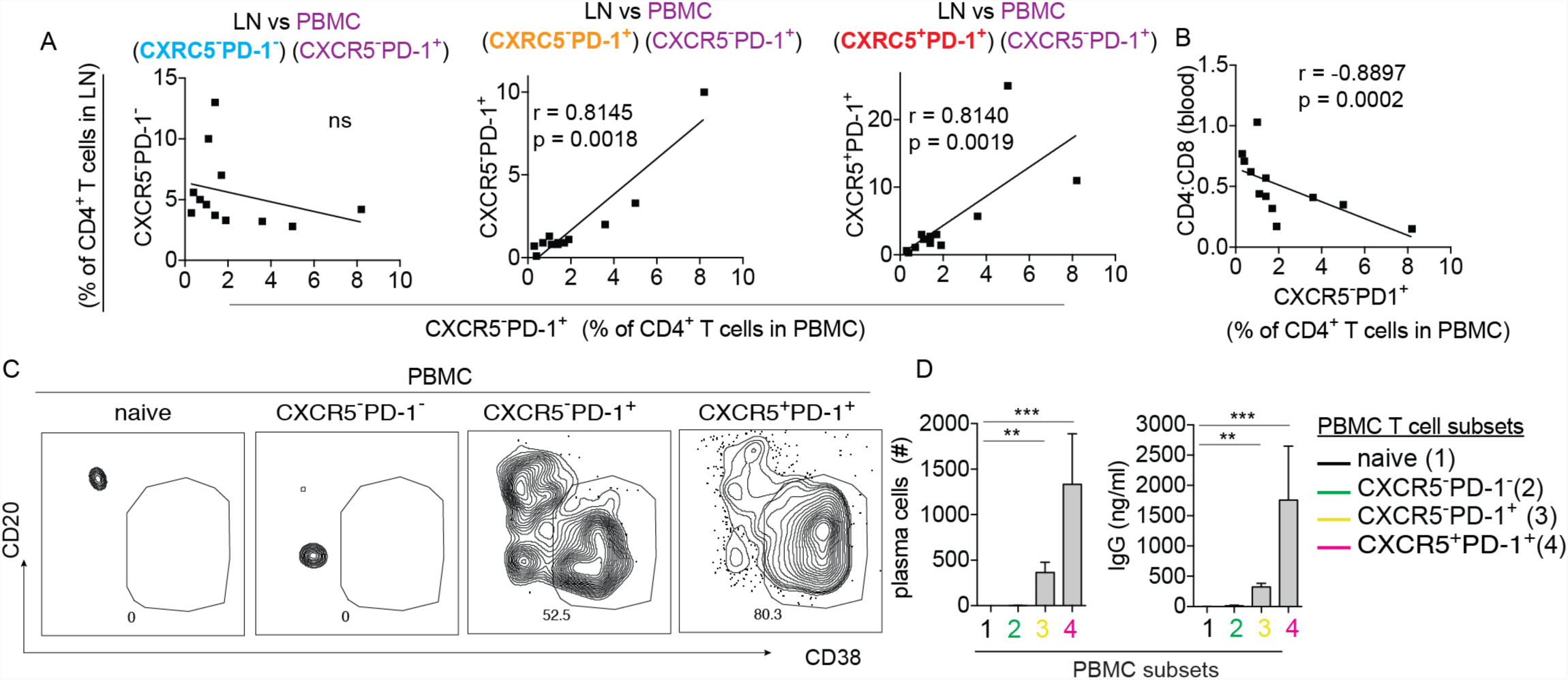
CXCR5^-^PD-1^+^ T cells in the blood reflect changes in the LN and promote B cell response in vitro. (A) Correlation between CXCR5^-^PD-1^+^ T cells in the PBMC versus different LN subsets (n = 12 paired PBMC and LNs) (B) Correlation between CXCR5^-^PD-1^+^ T cells in the PBMC and CD4:CD8 ratio. Data are from 12 HIV^+^ patients. Also see Fig. S2C-D. (C) Sorted CD4^+^ T cell subsets from the PBMCs of HIV infected individuals were cultured with memory B cells in the presence or absence of SEB for 7 days. Plots show representative staining for plasma cells in cocultures. (D) Bar-graph quantifies plasma cell and IgG level on day 7 of B cells cocultured with naïve cells (n = 8), CXCR5^-^PD-1^-^ T cells (n = 8), CXCR5^-^PD-1^+^ T cells (n = 7), or CXCR5^+^PD-1^+^ T cells (n = 5). Cocultures were performed in duplicates whenever possible with the available numbers of cells. For (A) and (B), association was measured by Spearman correlation. For (D), Kruskal-Wallis test was performed and corrected using Dunn’s multiple comparisons test.

To evaluate functional similarity between CXCR5^-^PD-1^+^ T cells in the PBMC and the lymphoid counterparts, we tested if CXCR5^-^PD-1^+^ T cells in the PBMCs are capable of driving B cell responses. We isolated naïve, CXCR5^-^PD-1^-^, CXCR5^-^PD-1^+^, or CXCR5^+^PD-1^+^ T cells from the PBMC and performed coculture assays with memory B cells in SEB. As expected, PBMC-derived CXCR5^+^PD-1^+^ T cells induced plasma cell differentiation and antibody production. In addition, we also observed induction of a lower but significant increase in plasma cell number and IgG level by CXCR5^-^PD-1^+^ T cells (Fig. 7C-D). Taken together, these data suggest that CXCR5^-^PD-1^+^ phenotype marks a migratory T cell population from the LN that contributed to the B cell help function of CXCR5^-^ T cells in the PBMCs.

## Discussion

T cell activation have been linked to rapid progression to AIDS prior to effective anti-viral therapy (Balagopal et al., 2015; Giorgi et al., 1993; Karim et al., 2013) and continues to be associated with an increase in non-AIDS related morbidities and mortalities in virally-suppressed patients (Hunt et al., 2016). While activated T cells have been known to accumulate in HIV Infected LNs (Biancotto et al., 2007), the phenotypic and functional diversity of activated T cells in lymphoid environment remains poorly defined. Here, we applied a complementary set of approaches on primary LNs obtained from a cohort of mostly untreated HIV^+^ patients to directly examine the phenotypic and functional diversity of activated CD3^+^ T cells in infected LNs. Consistent with the literature, HIV infected LNs are enriched for activated T cells that express the typical T_FH_ cell markers, CXCR5 and PD-1 (Lindqvist et al., 2012; Matthieu et al., 2013). In addition, we identified an accumulation of activated CXCR5 negative subset marked by high PD-1 expression in HIV infected LNs.

According to the prevailing model of T_FH_ cell differentiation, CXCR5 is expressed in early T_FH_ cells and it remains expressed during progressive stages of T_FH_ cell differentiation (Crotty, 2014). CD4^+^ T cells in the LNs that lacked CXCR5 expression have generally been categorized as a distinct population belonging to non-follicular CD4^+^ lineages. The focus on CXCR5 as a key T_FH_ cell marker is explained by its critical role in guiding T cells to the B cell zone to promote B cell differentiation and affinity maturation (Arnold et al., 2007; Haynes et al., 2007). However, T_FH_ cells may not be in continuous contact with B cells. What becomes of a T_FH_ cell over time remains unclear. Existing evidence suggest that T_FH_ cells can acquire a memory phenotype and circulate in the blood as CXCR5^+^ T cells (Bentebibel et al., 2013; Chevalier et al., 2011; He et al., 2013; Locci et al., 2013; Morita et al., 2011). Here, we provided evidence for an alternative phenotypic state under chronic inflammation that leads to the generation of CXCR5^-^ T cells.

We had initially hypothesized that CXCR5^-^PD-1^+^ T cells in HIV infected LNs reflected T cell exhaustion from chronic antigen stimulation. While CXCR5^-^PD-1^+^ T cells indeed expressed higher levels of exhaustion-related surface markers and RNA transcripts, these T cells retained the ability to produce cytokines and were functionally competent to promote plasma cell differentiation and antibody production in coculture assays. Notably, *in vitro* stimulated CXCR5^-^PD-1^+^ T cells produced an abundance of IFN-γ. IFN-γ has been shown to drive B cell expression of T-bet in the presence of TLR7 and 9 agonists (Naradikian et al., 2016). These T-bet^+^ B cells are typically found at a low frequency but increase with age and chronic inflammation (Hao et al., 2011; Moir et al., 2008; Rubtsova et al., 2013), and importantly, make up a major component of HIV-specific B cell responses in the setting of HIV infection (Knox et al., 2017; Moir et al., 2008). How B cells acquire T-bet expression in inflamed LNs remains unresolved. While our data do not directly demonstrate functional activity of CXCR5^-^PD-1^+^ T cells in the *in vivo* setting due to the limitation of our experimental system, the in vitro activity of CXCR5^-^PD-1^+^ T cells in coculture assays, along with correlative changes in T-bet^+^ B cells, suggest a role for CXCR5^-^PD-1^+^ T cells as potential regulators of B cell responses in inflamed LNs.

Where do these CXCR5^-^PD-1^+^ T cells come from? Several lines of evidence suggested that CXCR5^-^PD-1^+^ and CXCR5^+^PD-1^+^T cells share a common clonal lineage. First, the TCR repertoire from CXCR5^-^PD-1^+^ T cells overlaps with that of the TCR repertoire of CXCR5^+^PD-1^+^ T cells, with identical TCRβ CDR3 sequences found in both CXCR5^+^ and CXCR5^-^ subsets; second, in vitro activation down regulated CXCR5 expression on dividing T_FH_ cells; and third, ATAC-seq analyses showed an accessible open chromatin state around CXCR5 promoter region for CXCR5^-^PD-1^+^ T cells, which globally mapped in close proximity to CXCR5^+^PD-1^+^ T cells. The parsimonious explanation for these observations is that activation-dependent down regulation of CXCR5 on T_FH_ cells led to the accumulation of CXCR5^-^PD-1^+^ T cells. However, our data does not rule out the possibility that there are additional processes that coexist with this model. Differences in the source of precursor cells and/or the chosen differentiation pathway may contribute to the heterogeneity of phenotypic subsets within CXCR5^-^PD-1^+^ T cells, which remains to be fully elucidated in future studies.

In agreement with flexibility of CXCR5 expression, over 60% of transferred antigen-specific CXCR5^+^ memory cells in a lymphocytic choriomeningitis virus infection model recalled a T_FH_ cell response, but the remaining cells became CXCR5^-^ effectors (Hale et al., 2013). In humans, CXCR5^-^CD4^+^ T cells had been noted to generate low background level of antibody production in some experimental settings (Matthieu et al., 2013) (Chevalier et al., 2011; Rimpei et al., 2011). Recently, Brenner and colleagues directly examined CXCR5^-^ CD4^+^ T cells in the synovial tissue and blood from patients with rheumatoid arthritis and identified a CXCR5^-^ PD-1^+^ population characterized by CCR2 and HLA-DR expression that promoted plasma cell differentiation and antibody production. Based on the abundance of these CXCR5^-^ T cells in synovial lymphoid aggregates and their proximity to B cells, Rao et al. suggested that infiltration by CXCR5^-^PD-1^+^ T cells may play a key role in the formation of tertiary lymphoid aggregates in chronically inflamed joints (Rao et al., 2017). Our data extends these findings to the lymphoid compartment by identifying a phenotypically similar CXCR5 negative population that also provided help to B cells in HIV infected LNs. The origin of CXCR5 negative atypical B cell helping T cells in the blood remains unknown as circulating CXCR5^+^ cells do not become CXCR5 negative *in vitro* (Rao et al., 2017). Our data suggest that CXCR5^-^PD-1^+^ T cells in the LN acquire migratory program and contribute to the emergence of atypical B cell helping activity in peripheral circulation. The CXCR5^-^PD-1^+^ T cells that we identified in HIV infected LNs are notable for a strong cell migration signals that included upregulation of numerous surface receptors involved in cell trafficking and LN egress. Movement of expanded CXCR5^-^PD-1^+^ T cells from the LN into blood could explain the overlap in shared TCRs between CXCR5^-^PD-1^+^ T cells in the lymphoid and circulatory compartments. As RA and HIV are both disease that involve chronic inflammation, the accumulation of a migratory T_FH_-like population in the LN may represent a general state of lymphoid dysregulation that contributes to inflammatory responses in peripheral compartments.

In summary, we have defined a CXCR5^-^PD-1^+^ T cell population with B cell help functionality that becomes expanded in HIV infected LNs and recognizes HIV antigens. The association between these cells with more severe clinical presentation and HIV-induced changes in B cell distribution highlights the importance of CXCR5^-^CD4^+^ T cells on B cell responses in HIV infection. Finally, the accumulation of CXCR5^-^PD-1^+^ T cells in lymphoid tissues and their impact on B cell responses are unlikely to be limited to chronic viral diseases. Insights into activated lymphoid environment will be broadly applicable for understanding immune dysregulation that occur in other states of chronic inflammation, including aging and autoimmunity.

## Acknowledgments

We thank Dr. Ke-Yue Ma for helping with TCR sequencing run. We also thank the NIH AIDS Reagent Program and Penn Center for AIDS Research (P30 AI 045008) for providing reagents and samples.

## Funding

VA Merit Award IMMA-020-15F (L.F.S), NIH R01AI134879 (L.F.S), R00AG040149 (N.J.), S10OD020072 (N.J.), the Welch Foundation grant F1785 (N.J.).

## Author Contributions

L.F.S. designed the study. D.D.A. performed CyTOF and FACS phenotypic analyses and coculture assays. Y. W performed RNA sequencing and ATAC-seq and assisted with data analyses. J. L performed the data analyses for RNA and ATAC-seq. C. H. and B.S.W., performed TCR data analysis. B.S.W, M.J.M. and S.M.H performed TCR sequencing experiment. P.D.R., Y.A.T., and G.R.T established the infrastructure for HIV^+^ patient recruitment and provided HIV infected LN samples and the associated clinical information. A.N. established the infrastructure for sample collection from organ transplant donors and provided the LN from healthy individuals. I. F. maintained the CFAR clinical database/ biorepository and provided a subset of PBMC samples from HIV^+^ individuals. L.F.S., D.D.A., and Y. W. wrote and edited the manuscript.

## Declaration of Interests

N.J. is a scientific advisor for ImmuDX LLC and Immune Arch Inc. Other authors do not have conflict of interest to declare.

## Contact for reagents and resource sharing

Correspondence and requests for materials should be addressed to Laura F. Su.

## Experimental methods and subject detail

### Study Design

LNs and PBMCs were obtained from HIV^+^ patients undergoing excision of palpable cervical LNs for clinical diagnostic workup in Mexico. Additional PBMCs were obtained from a second cohort of patients at the University of Pennsylvania. LNs from healthy controls were obtained from organ transplant donors. Sample sizes were not pre-specified and were dictated by the availability of the samples, which were collected over four years. All samples were de-identified and obtained with IRB regulatory approval from the University of Pennsylvania. Recruitment of HIV^+^ Mexican study subjects was additionally approved by the Comité de Ciencia y de Ética en Investigación and the Comité de Investigación of the Instituto Nacional de Enfermedades Respiratorias (INER) in México city. Subject characteristics are shown in Table S1.

### CyTOF staining and data analyses

Cryopreserved cells were thawed in benzonase (0.025 unit/ml, Sigma) and stained with metal-conjugated antibody panel (Table S2). T cell stimulation was performed by a 5-hour incubation in PMA (5ng/ml, Sigma) and ionomycin (50ng/ml, Sigma) in the presence monensin (2mM, Sigma) and Brefeldin A (5ug/ml, Sigma) as previously described (Wendel et al., 2018). Antibody stained cells were mixed with normalization beads and acquired on CyTOF 2. Bead standards were used to normalize CyTOF runs with the Matlab-based Nolan lab normalizer (Finck et al., 2013). Iridium+Cisplatin-cells were excluded for doublets and beads. Equal numbers of manually gated CD38^+^HLA-DR^+^ and CD38-HLA-DR-T cells were downsampled for each sample and exported using FlowJo. Dimensionality reduction, subset identification, and progression for these populations were performed using the “R” package, “cytofkit” in a single batch. Dimensionality reduction was performed using t-SNE (n=15979, perplexity=30, iteration=1000, theta=0.5). The subsequent cluster identification of these populations was performed using cytofkit’s Phenograph (k=15) implementation. (cluster markers: CD25, IL2, ICOS, Ki67, TCF1, PD1, CCR7, CD45RO, CCR5, CD71, IL-4, Foxp3, CD57, perforin. CD5, granzymeB, IFN-γ, CD69, TNF-α, CD27, CXCR4, CXCR5, CD103, CCR4, CCR6, IL-13, IL-17A). The diffusion maps of the Phenograph clusters were created after downsampling each cluster to 500 cells, using euclidean distance calculation with an output dimensionality of 4. The heatmap was generated using the “gplot” package in R and show the raw staining intensity of each marker after arcsinh transformation. Heatmap dendograms were clustered by Euclidean distance.

### Cell sorting and staining by flow cytometry

Cryopreserved cells were thawed in 0.025 unit/mL of benzonase (Sigma). For surface staining, cells were stained with antibody cocktails for 30 minutes at room temperature. Fixable aqua dye (ThermoFisher) staining was used for live/dead discrimination. T cell stimulation was performed as described above by adding PMA and ionomycin for 5 hours in the presence monensin and Brefeldin A. For intracellular cytokine or transcription factor staining, cells were permeabilized and fixed using Foxp3 staining buffer set and stained in permeabilization buffer according to vendor protocol (eBioscience). Samples were acquired on LSRII (BD Biosciences). For cell sorting, freshly thawed T cells were separated into ICOS high or ICOS low subset by positive enrichment using a magnetic column (Miltenyi Biotec). Figure S6A shows gating strategy for sorting CXCR5^+^PD-1^+^, CXCR5^-^PD-1^+^, CXCR5^-^PD-1^-^, and naïve subpopulations for TCR sequencing, RNA sequencing, ATAC-seq, and coculture assays (Fig. S6A). ICOS staining enriched for activated CXCR5^-^PD-1^+^ T cells (Fig. S6B). CD27 stained the majority CXCR5^-^PD-1^+^ T cells from samples used for RNA sequencing and ATAC-seq (Fig. S6C). For PBMC, PD-1^+^ICOS^+^ gate was set by the absence of double positive T cells in the naïve population as indicated (Fig. S6D). Memory B cells were sorted as live CD19^+^CD3^-^CD4^-^IgD^-^CD38^-^ cells (Fig. S6A). Cell sorting was performed using FACSAria III (BD Biosciences). Analyses were performed using FlowJo 10.0.8 (Tree Star).

### T:B coculture assays and antigen-specific analyses

Memory B cells (5,000 cells/well) were cultured with T cells (5,000 cells/well) in a V-bottom plate with media supplemented with the anti-retroviral drugs Raltegravir (1uM, Cayman Chemical) and Efavorez (1uM, Cayman Chemical). Cells were cultured in RPMI (Mediatech) and stimulated with DMSO or SEB (1ug/ml, Toxin Technology). Total IgG was measured in a standard ELISA assay using 96-well medium binding plates (Greiner) coated with IgG F(ab’) 2GT antibody (1:100, Bethyl) and incubated with serial dilutions of supernatants and IgG standards (Sigma-Aldrich). Plate bound IgG was detected using biotinylated anti-IgG (1:2000, BD) and streptavidin-HRP (1:2000, Southern Biotech). Plasma cells were identified as live CD19^+^CD4^-^CD20^-^CD38^+^ B cells. For peptide stimulation, cells were incubated with 0.25ug/ml of Gag and Env peptide pools for 18 hours (#12425 and #12540, NIH AIDS Reagent Program). Peptide-reactive T cells were identified by OX40 and CD25 expression. For LN samples, sorted T cells were stained with CFSE (3uM, Invitrogen) to identify dividing T cells. Measurements of IgG level, B and T cell phenotypes were assayed after 7 days in culture.

### TCRβ sequencing and analyses

TCR library generation and sequencing on FACS-sorted T cells was performed as previously described (Wendel et al., 2018). Briefly, FACS-sorted cells were lysed, and total RNA was purified using AllPrep DNA/RNA Micro Kit (Qiagen). 30% of the purified RNA was used for library generation and sequencing. Consensus sequences were constructed within each molecular identifier (MID) group (Ke-Yue Ma, 2018). Consensus TCR sequences were subjected to the CDR3 blast module of MIGEC (Shugay et al., 2014) to assign V and J alleles and parse out the CDR3 sequence (Table S3). Bhattacharyya coefficient was used to measure TCR repertoire similarity between CXCR5^-^PD-1^+^ T cells with other populations within LN and between CXCR5^-^PD-1^+^ T cells in the PBMC and LN subsets (Bhattacharyya, 1943). Circos plots were generated using circlize R package to visualize clones shared between T cell subsets.

### ATAC-seq and analyses

ATAC-seq was performed as previously described with minor modifications (Buenrostro et al., 2013). In brief, sorted T cells were pelleted and resuspended in 50uL lysis buffer (10 mM Tris-HCl [pH 7.4], 10 mM NaCl,3 mM MgCl2, 0.1% IGEPAL CA-630). Transposition reaction was carried out immediately in a 25ul reaction with 1.25ul of Tn5 transposase (Illumina) in a 37oC water bath for 37 minutes to tag and fragment accessible chromatin. Tagmented DNA was purified using a MinElute Reaction Cleanup Kit (QIAGEN) and amplified with 12 cycles of PCR. Sample quality was determined using a Tapestation 2200 system (Agilent). Libraries were purified using Agencourt AMPure XP kit (Axygen). Library concentrations were determined using Qubit dsDNA HS Assay Kit (Invitrogen). Libraries were paired-end sequenced on a NextSeq 550 (Illumina). All the reads are quality-trimmed with trimmomatic-0.36-5 (see Table S3). The paired-end 38bp reads were aligned to human assembly hg19 using bowtie alignment tool (bowtie2-2.3.4.1). The peaks were called using callpeaks function in MACS2. Peaks called with a q-value cutoff of 0.05 (Benjamin-Hochberg correction) were analyzed with DiffBind R package. The intervals for peak comparison were re-centered around the most enriched points with a 500-bp window. The consensus peaksets were then normalized and used as input for differential analysis by DESeq2 with a significance cutoff FDR at 0.05 (Benjamin-Hochberg correction). DiffBind was used to generate the PCA plot.

### RNA sequencing and analyses

Sorted T cells were pelleted and resuspended in Buffer RLT Plus (QIAGEN) with 10% 2-Mercaptoethanol (Sigma). Total RNA from each sample was isolated using the RNeasy Plus Micro Kit (QIAGEN). RNA integrity was determined using a 2100 Bioanalyzer (Agilent). Reverse transcription and cDNA synthesis was performed using the SMART-Seq v4 Ultra Low Input RNA Kit for Sequencing (Clontech). cDNA concentrations were determined using Qubit dsDNA HS Assay Kit (Invitrogen). DNA libraries were prepared using Nextera XT Library Prep Kit (Illumina) and sequenced on NextSeq 550 (Illumina). All the reads are quality-trimmed with trimmomatic-0.36-5 (see Table S3). The paired-end 38bp reads were aligned to human assembly using STAR alignment tool (STAR 2.6). For finding differential expressed genes between samples, DESeq2 R packaged was used to calculate variation, with a significance cutoff padj at 0.05 (Benjamin-Hochberg correction). PCA analysis was performed using plotPCA function in DESeq2 R package. The variant genes were then submitted to GOrilla for gene ontology analysis. Gene Set Enrichment Analysis (GSEA) was performed to compare the enriched genes in CXCR5^+^PD-1^+^ / CXCR5^-^PD-1^+^ T cells. The significant hit from GSEA and Gorilla were matched to generate a list of enriched gene ontologies shared by both methods. Supervise hierarchical clustering heatmap for differentially expressed genes that contributed to the GO terms was generated gplots R package, using scaled, normalized expression values as input.

## Quantification and statistical analysis

Assessment of normality was performed using D’Agostino-Pearson test. Pearson or Spearman correlation was used depending on the normality of the data to measure the degree of association. The best-fitting line was calculated using least squares fit regression. Statistical comparisons were performed using two-tailed Student’s t-test or Wilcoxon signed-rank test, using a p-value of <0.05 as a cutoff to determine statistical significance. Multiple-way comparisons were performed using ANOVA or nonparametric tests (Friedman test for matched measures or Kruskal-Wallis test for unmatched measures). Statistical analyses were performed using GraphPad Prism. * *P* < 0.05; ** *P* < 0.005; *** *P* < 0.0005. **** *P* < 0.0001.

## Supplementary Information

**Table S1:**
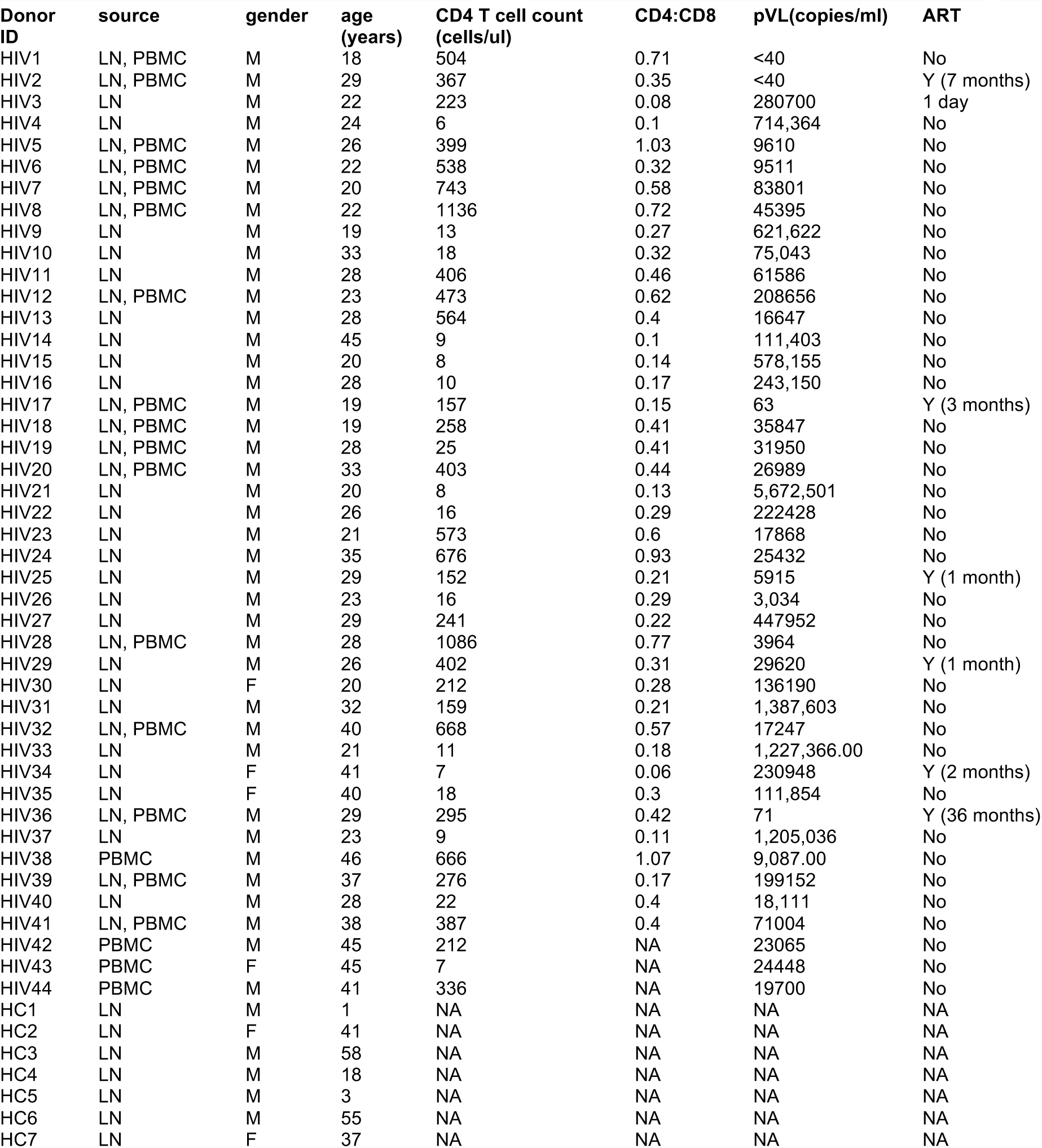
Clinical characteristics and demographic information of LN and PBMC samples

| Donor ID | source | gender | age (years) | CD4 T cell count (cells/ul) | CD4:CD8 | pVL(copies/ml) | ART |
| --- | --- | --- | --- | --- | --- | --- | --- |
| HIV1 | LN, PBMC | M | 18 | 504 | 0.71 | <40 | No |
| HIV2 | LN, PBMC | M | 29 | 367 | 0.35 | <40 | Y (7 months) |
| HIV3 | LN | M | 22 | 223 | 0.08 | 280700 | 1 day |
| HIV4 | LN | M | 24 | 6 | 0.1 | 714,364 | No |
| HIV5 | LN, PBMC | M | 26 | 399 | 1.03 | 9610 | No |
| HIV6 | LN, PBMC | M | 22 | 538 | 0.32 | 9511 | No |
| HIV7 | LN, PBMC | M | 20 | 743 | 0.58 | 83801 | No |
| HIV8 | LN, PBMC | M | 22 | 1136 | 0.72 | 45395 | No |
| HIV9 | LN | M | 19 | 13 | 0.27 | 621,622 | No |
| HIV10 | LN | M | 33 | 18 | 0.32 | 75,043 | No |
| HIV11 | LN | M | 28 | 406 | 0.46 | 61586 | No |
| HIV12 | LN, PBMC | M | 23 | 473 | 0.62 | 208656 | No |
| HIV13 | LN | M | 28 | 564 | 0.4 | 16647 | No |
| HIV14 | LN | M | 45 | 9 | 0.1 | 111,403 | No |
| HIV15 | LN | M | 20 | 8 | 0.14 | 578,155 | No |
| HIV16 | LN | M | 28 | 10 | 0.17 | 243,150 | No |
| HIV17 | LN, PBMC | M | 19 | 157 | 0.15 | 63 | Y (3 months) |
| HIV18 | LN, PBMC | M | 19 | 258 | 0.41 | 35847 | No |
| HIV19 | LN, PBMC | M | 28 | 25 | 0.41 | 31950 | No |
| HIV20 | LN, PBMC | M | 33 | 403 | 0.44 | 26989 | No |
| HIV21 | LN | M | 20 | 8 | 0.13 | 5,672,501 | No |
| HIV22 | LN | M | 26 | 16 | 0.29 | 222428 | No |
| HIV23 | LN | M | 21 | 573 | 0.6 | 17868 | No |
| HIV24 | LN | M | 35 | 676 | 0.93 | 25432 | No |
| HIV25 | LN | M | 29 | 152 | 0.21 | 5915 | Y (1 month) |
| HIV26 | LN | M | 23 | 16 | 0.29 | 3,034 | No |
| HIV27 | LN | M | 29 | 241 | 0.22 | 447952 | No |
| HIV28 | LN, PBMC | M | 28 | 1086 | 0.77 | 3964 | No |
| HIV29 | LN | M | 26 | 402 | 0.31 | 29620 | Y (1 month) |
| HIV30 | LN | F | 20 | 212 | 0.28 | 136190 | No |
| HIV31 | LN | M | 32 | 159 | 0.21 | 1,387,603 | No |
| HIV32 | LN, PBMC | M | 40 | 668 | 0.57 | 17247 | No |
| HIV33 | LN | M | 21 | 11 | 0.18 | 1,227,366.00 | No |
| HIV34 | LN | F | 41 | 7 | 0.06 | 230948 | Y (2 months) |
| HIV35 | LN | F | 40 | 18 | 0.3 | 111,854 | No |
| HIV36 | LN, PBMC | M | 29 | 295 | 0.42 | 71 | Y (36 months) |
| HIV37 | LN | M | 23 | 9 | 0.11 | 1,205,036 | No |
| HIV38 | PBMC | M | 46 | 666 | 1.07 | 9,087.00 | No |
| HIV39 | LN, PBMC | M | 37 | 276 | 0.17 | 199152 | No |
| HIV40 | LN | M | 28 | 22 | 0.4 | 18,111 | No |
| HIV41 | LN, PBMC | M | 38 | 387 | 0.4 | 71004 | No |
| HIV42 | PBMC | M | 45 | 212 | NA | 23065 | No |
| HIV43 | PBMC | F | 45 | 7 | NA | 24448 | No |
| HIV44 | PBMC | M | 41 | 336 | NA | 19700 | No |
| HC1 | LN | M | 1 | NA | NA | NA | NA |
| HC2 | LN | F | 41 | NA | NA | NA | NA |
| HC3 | LN | M | 58 | NA | NA | NA | NA |
| HC4 | LN | M | 18 | NA | NA | NA | NA |
| HC5 | LN | M | 3 | NA | NA | NA | NA |
| HC6 | LN | M | 55 | NA | NA | NA | NA |
| HC7 | LN | F | 37 | NA | NA | NA | NA |

**Table S2:** CyTOF antibody panels

Panel 1
| Antibody | Metal | Clone | Vendor |
| --- | --- | --- | --- |
| CD57 | 113 | HCD57 | Biolegend |
| CD3 | 141 | UCHT1 | BD |
| CD5 | 142 | UCHT2 | Biolegend |
| CD8 | 143 | SK1 | Biolegend |
| CD4 | 144 | SK3 | Biolegend |
| CD19 | 145 | HIB19 | Biolegend |
| Granzyme B | 146 | CLB-GB11 | eBioscience |
| IFN-γ | 147 | 4S.B3 | eBioscience |
| HLA-DR | 148 | L243 | biolegend |
| CD14 | 149 | M5E2 | Bio Legend |
| CD69 | 150 | FN50 | Biolegend |
| CD38 | 151 | HB-7 | Biolegend |
| TNF-α | 152 | MAB11 | biolegend |
| CD45RO | 153 | UCHL1 | biolegend |
| CD27 | 154 | LG.7F9 | eBioscience |
| TCR$\alpha\beta$ | 155 | T10B9.1A-31 | BD |
| CCR5 | 156 | J418F1 | Biolegend |
|  | 157 |  |  |
| CD71 | 158 | CY1G4 | biolegend |
| CXCR4 | 159 | 12G5 | biolegend |
| IL-4 | 160 | 8D4-8 | BD |
| CD25 | 161 | M-A251 | BD |
| IL-2 | 162 | MQ1-17H12 | biolegend |
| ICOS | 163 | c398-4A | biolegend |
| Ki67 | 164 | B56 | BD |
| Foxp3 | 165 | PCH101 | ebiosciences |
| TCF1 | 166 | C63D9 | Cell Signaling |
| PD-1 | 167 | EH12.2H7 | Biolegend |
| CCR7 | 168 | G043H7 | biolegend |
| CXCR5 | 169 | RF8B2 | BD |
| TCR$\gamma\delta$ | 170 | SA6.E9 | invitrogen |
| CD103 | 171 | B-Ly7 | ebioscience |
| CCR4 | 172 | 1G1 | BD |
| CCR6 | 173 | G034E3 | biolegend |
| IL-13 | 174 | JES10-5A2 | biolegend |
| Perforin | 175 | DG9 | biolegend |
| IL-17A | 176 | BL168 | Biolegend |

Panel 2 (Wendel et al., 2018)
| antibodies | metal | clone | Vendor |
| --- | --- | --- | --- |
| CD57 | 113 | HCD57 | Biolegend |
| CD3 | 141 | UCHT1 | BD |
| CD5 | 142 | UCHT2 | Biolegend |
| CD8 | 143 | SK1 | Biolegend |
| CD4 | 144 | SK3 | Biolegend |
| CD19 | 145 | HIB19 | Biolegend |
| IgD | 146 | IA6-2 | Biolegend |
| IFN-γ | 147 | 4S.B3 | eBioscience |
| CCR6 | 148 | G034E3 | Biolegend |
| CD14 | 149 | M5E2 | Biolegend |
| CD69 | 150 | FN50 | Biolegend |
| Granzyme A | 151 | CB9 | Biolegend |
| TNF-α | 152 | MAB11 | Biolegend |
| CD45RO | 153 | UCHL1 | Biolegend |
| CD27 | 154 | LG.7F9 | eBioscience |
| TCRab | 155 | T10B9.1A-31 | BD |
| CCR5 | 156 | J418F1 | Biolegend |
| Ki67 | 157 | B56 | BD |
| BLYS | 158 | 1D6 | Biolegend |
| BCL6 | 159 | K112-91 | BD |
| IL-4 | 160 | 8D4-8 | BD |
| CD25 | 161 | M-A251 | BD |
| Eomes | 162 | wd1928 | eBioscience |
| ICOS | 163 | c398-4A | Biolegend |
| IL-5 | 164 | TRFK5 | Biolegend |
| Foxp3 | 165 | PCH101 | eBioscience |
| CD95 | 166 | DX2 | Biolegend |
| CD45RA | 167 | HI100 | eBioscience |
| CCR7 | 168 | G043H7 | Biolegend |
| CXCR5 | 169 | RF8B2 | BD |
| Tbet | 170 | 4B10 | Biolegend |
| PD-1 | 171 | EH12.2H7 | Biolegend |
| IL-21 | 172 | 3A3-N2 | Biolegend |
| IL-2 | 173 | MQ1-17H12 | eBioscience |
| CD24 | 174 | ML5 | Biolegend |
| CXCR3 | 175 | G025H7 | Biolegend |
| CD38 | 176 | HIT2 | Biolegend |

**Table S3:** Cell and transcript counts for sequencing experiments

| Donor ID | Assay Type | T cell Population | site | cell count | mapped reads after QC | Donor ID | Assay Type | T cell Population | site | cell count | transcript count |
| --- | --- | --- | --- | --- | --- | --- | --- | --- | --- | --- | --- |
| HIV4 | ATAC-seq | CXCR5-PD-1- | LN | 10000 | 49048896 | HIV13 | TCR-seq | CXCR5-PD-1- | LN | 10000 | 6533 |
| HIV16 | ATAC-seq | CXCR5-PD-1- | LN | 17,544 | 52354592 | HIV28 | TCR-seq | CXCR5-PD-1- | LN | 10000 | 2821 |
| HIV21 | ATAC-seq | CXCR5-PD-1- | LN | 20,000 | 53542641 | HIV41 | TCR-seq | CXCR5-PD-1- | LN | 10000 | 6254 |
| HIV4 | ATAC-seq | CXCR5-PD-1+ | LN | 10000 | 40979805 | HIV2 | TCR-seq | CXCR5-PD-1- | LN | 10000 | 8777 |
| HIV16 | ATAC-seq | CXCR5-PD-1+ | LN | 20,000 | 51161176 | HIV17 | TCR-seq | CXCR5-PD-1- | LN | 10000 | 8945 |
| HIV21 | ATAC-seq | CXCR5-PD-1+ | LN | 20,000 | 47122262 | HIV39 | TCR-seq | CXCR5-PD-1- | LN | 10000 | 9033 |
| HIV4 | ATAC-seq | CXCR5+P D-1+ | LN | 3122 | 44084338 | HIV13 | TCR-seq | CXCR5-PD-1+ | LN | 2500 | 4546 |
| HIV16 | ATAC-seq | CXCR5+P D-1+ | LN | 20,000 | 51623107 | HIV28 | TCR-seq | CXCR5-PD-1+ | LN | 2515 | 2708 |
| HIV21 | ATAC-seq | CXCR5+P D-1+ | LN | 20,000 | 51243858 | HIV41 | TCR-seq | CXCR5-PD-1+ | LN | 10000 | 10290 |
| HIV4 | ATAC-seq | naïve | LN | 10000 | 41241783 | HIV2 | TCR-seq | CXCR5-PD-1+ | LN | 5000 | 3182 |
| HIV16 | ATAC-seq | naïve | LN | 20,000 | 50541268 | HIV17 | TCR-seq | CXCR5-PD-1+ | LN | 2500 | 2836 |
| HIV21 | ATAC-seq | naïve | LN | 20,000 | 43842949 | HIV39 | TCR-seq | CXCR5-PD-1+ | LN | 4085 | 3542 |
|  |  |  |  |  |  | HIV13 | TCR-seq | CXCR5+P D-1+ | LN | 10000 | 17659 |
| HIV4 | RNA-seq | CXCR5-PD-1- | LN | 9500 | 55493810 | HIV28 | TCR-seq | CXCR5+P D-1+ | LN | 2500 | 4135 |
| HIV16 | RNA-seq | CXCR5-PD-1- | LN | 5000 | 56601410 | HIV41 | TCR-seq | CXCR5+P D-1+ | LN | 10000 | 10167 |
| HIV21 | RNA-seq | CXCR5-PD-1- | LN | 5000 | 63020800 | HIV2 | TCR-seq | CXCR5+P D-1+ | LN | 10000 | 10900 |
| HIV4 | RNA-seq | CXCR5-PD-1+ | LN | 33198 | 46402370 | HIV17 | TCR-seq | CXCR5+P D-1+ | LN | 8758 | 8936 |
| HIV16 | RNA-seq | CXCR5-PD-1+ | LN | 5000 | 56568020 | HIV39 | TCR-seq | CXCR5+P D-1+ | LN | 5000 | 19342 |
| HIV21 | RNA-seq | CXCR5-PD-1+ | LN | 5000 | 60278910 | HIV13 | TCR-seq | naïve | LN | 10000 | 18376 |
| HIV4 | RNA-seq | CXCR5+P D-1+ | LN | 6105 | 44762968 | HIV28 | TCR-seq | naïve | LN | 10000 | 5488 |
| HIV16 | RNA-seq | CXCR5+P D-1+ | LN | 5000 | 54385182 | HIV41 | TCR-seq | naïve | LN | 10000 | 5822 |
| HIV21 | RNA-seq | CXCR5+P D-1+ | LN | 5000 | 56737602 | HIV2 | TCR-seq | naïve | LN | 5000 | 3502 |
| HIV4 | RNA-seq | naïve | LN | 20000 | 48161000 | HIV17 | TCR-seq | naïve | LN | 10000 | 6750 |
| HIV16 | RNA-seq | naïve | LN | 5000 | 49000230 | HIV39 | TCR-seq | naïve | LN | 10000 | 4314 |
| HIV21 | RNA-seq | naïve | LN | 5000 | 54953878 | HIV13 | TCR-seq | CXCR5-PD-1+ | PBMC | 2500 | 3374 |
|  |  |  |  |  |  | HIV28 | TCR-seq | CXCR5-PD-1+ | PBMC | 2267 | 3016 |
|  |  |  |  |  |  | HIV41 | TCR-seq | CXCR5-PD-1+ | PBMC | 5000 | 6640 |
|  |  |  |  |  |  | HIV2 | TCR-seq | CXCR5-PD-1+ | PBMC | 2500 | 1711 |
|  |  |  |  |  |  | HIV17 | TCR-seq | CXCR5-PD-1+ | PBMC | 293 | 496 |
|  |  |  |  |  |  | HIV39 | TCR-seq | CXCR5-PD-1+ | PBMC | 4487 | 4536 |

**Figure S1:**
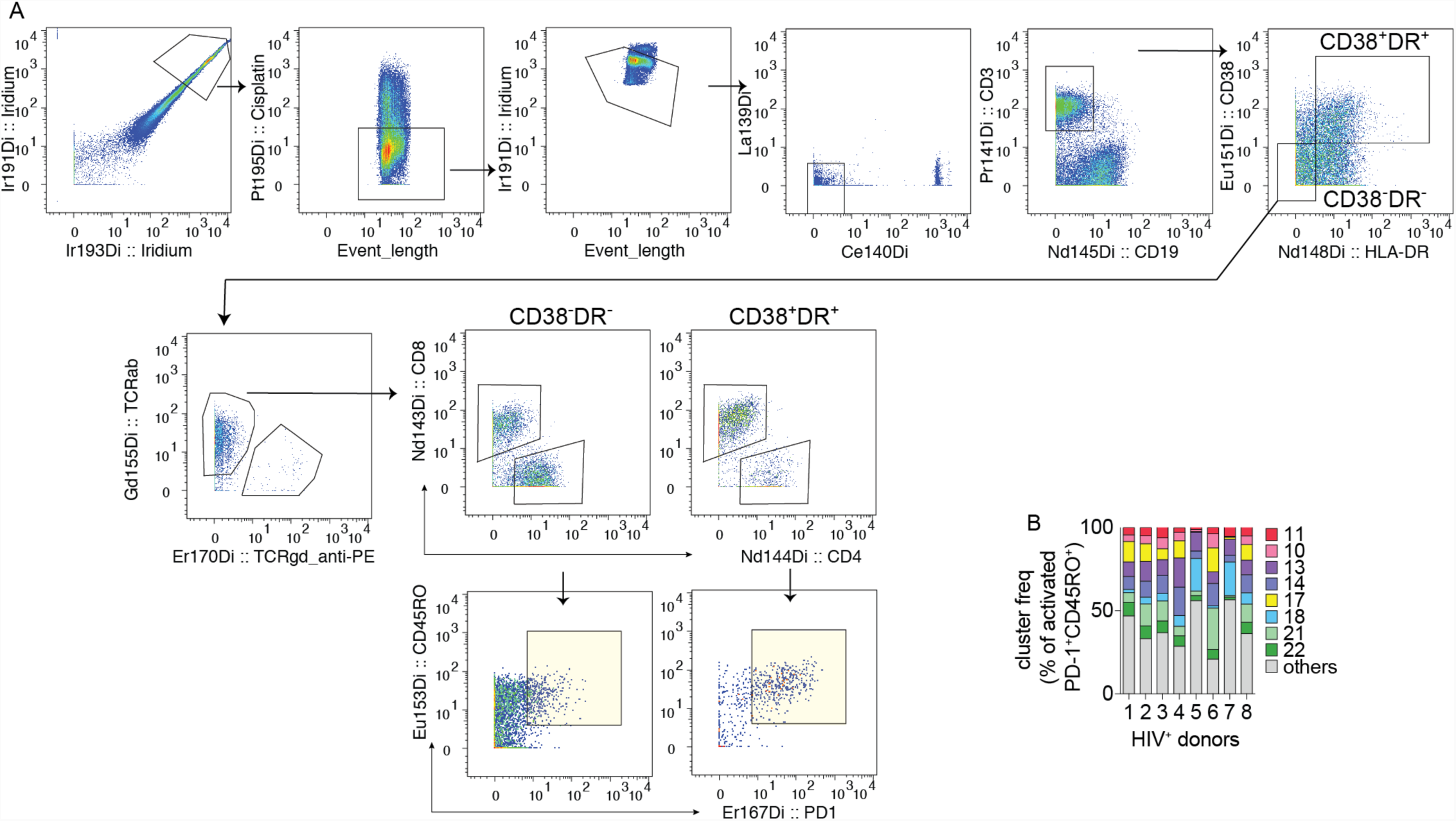
Identification of activated and quiescent T cells in LN samples by CyTOF. (A) Representative plots showing gating strategy for identifying the input cells used for CyTOF analyses (colored in yellow). (B) Bar-graph shows the relative frequency of the eight activation-enriched clusters within the CD38^+^HLA-DR^+^ population. Each bar represents one donor.

**Figure S2:**
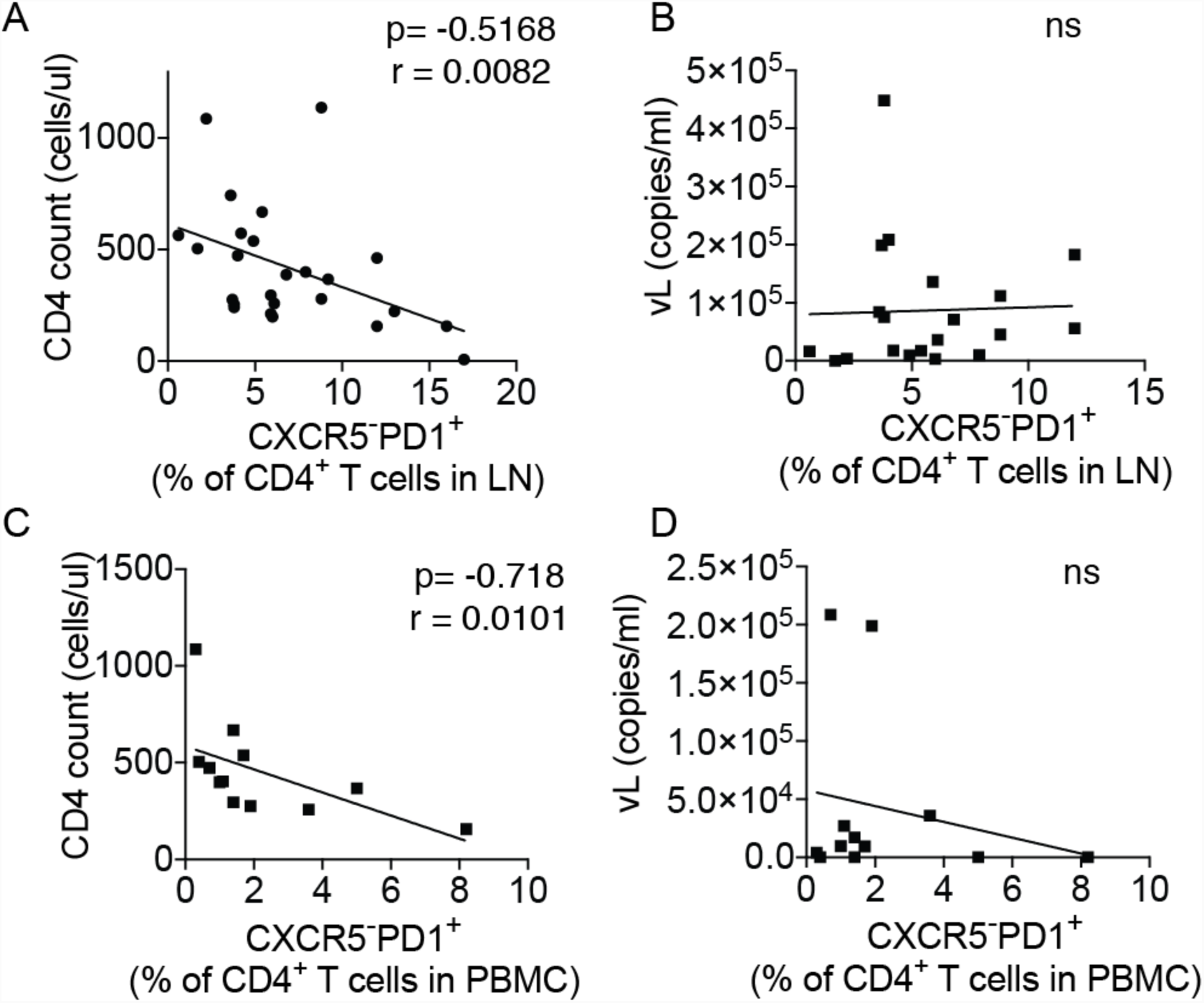
Correlation between CXCR5-PD-1^+^ T cell frequency and clinical parameters of HIV infection. (A-B) The frequency of CXCR5-PD-1^+^ T cells in the LN is associated with CD4^+^ T cell count (A) but not with viral load (B). Data include 25 HIV^+^ LN samples. (C-D) The frequency of PBMC-derived CXCR5-PD-1^+^ T cells is associated with CD4^+^ T cell count (A) but not with viral load (B). Data include 12 HIV^+^ PBMC samples. Association was measured by Spearman correlation and the best-fitting line was calculated using least squares fit regression.

**Figure S3:**
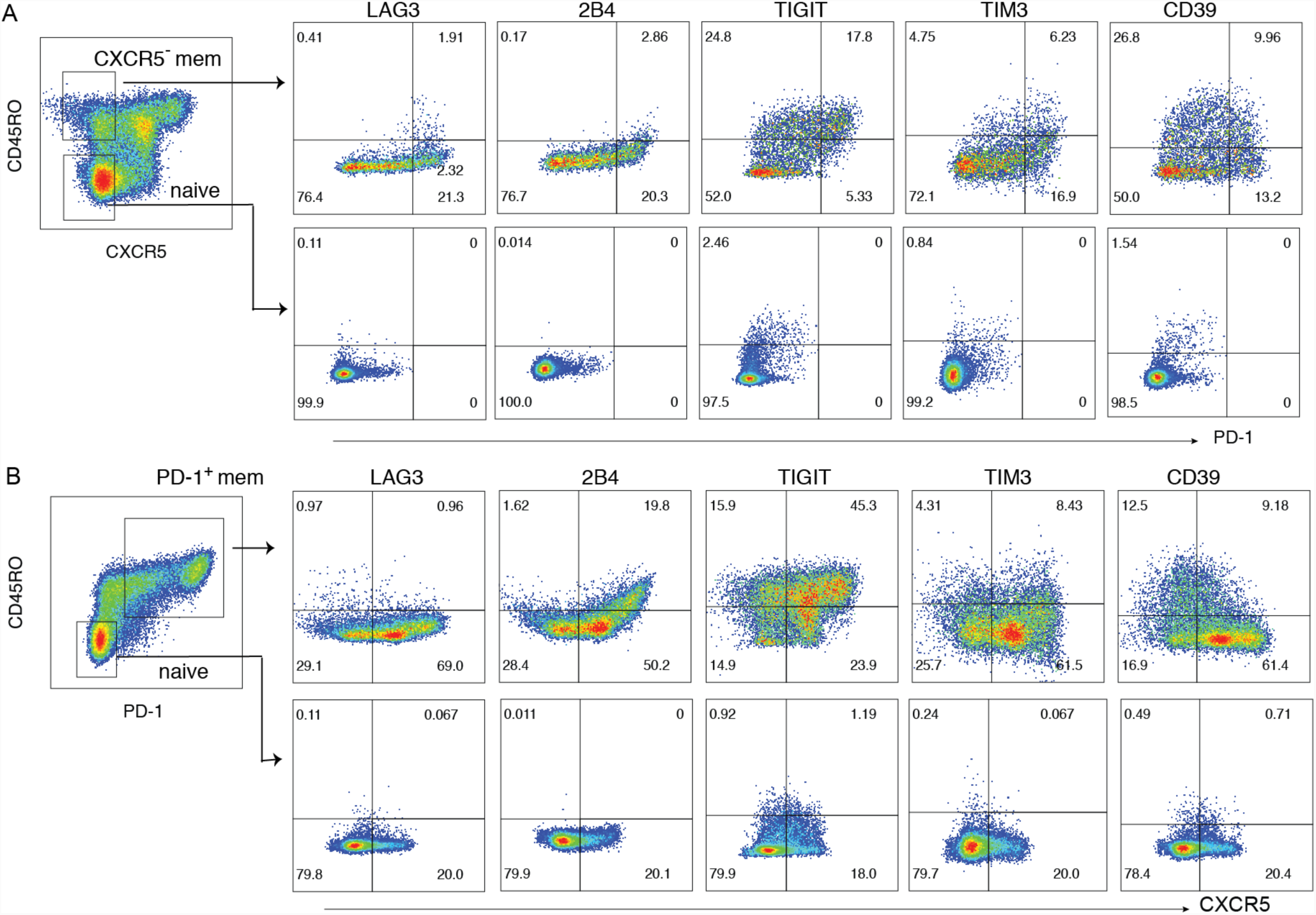
Expression of exhaustion-related surface molecules on CD4^+^ T cells in LNs. (A) Representative plots showing expression of LAG3, 2B4, TIGIT, TIM3, and CD39 on memory CD4^+^ T cells. CXCR5^+^ cells were excluded. (B) Representative plots showing expression of LAG3, 2B4, TIGIT, TIM3, and CD39 on PD-1^+^ memory T cells. Identical gates on naïve cells are shown for comparison.

**Figure S4:**
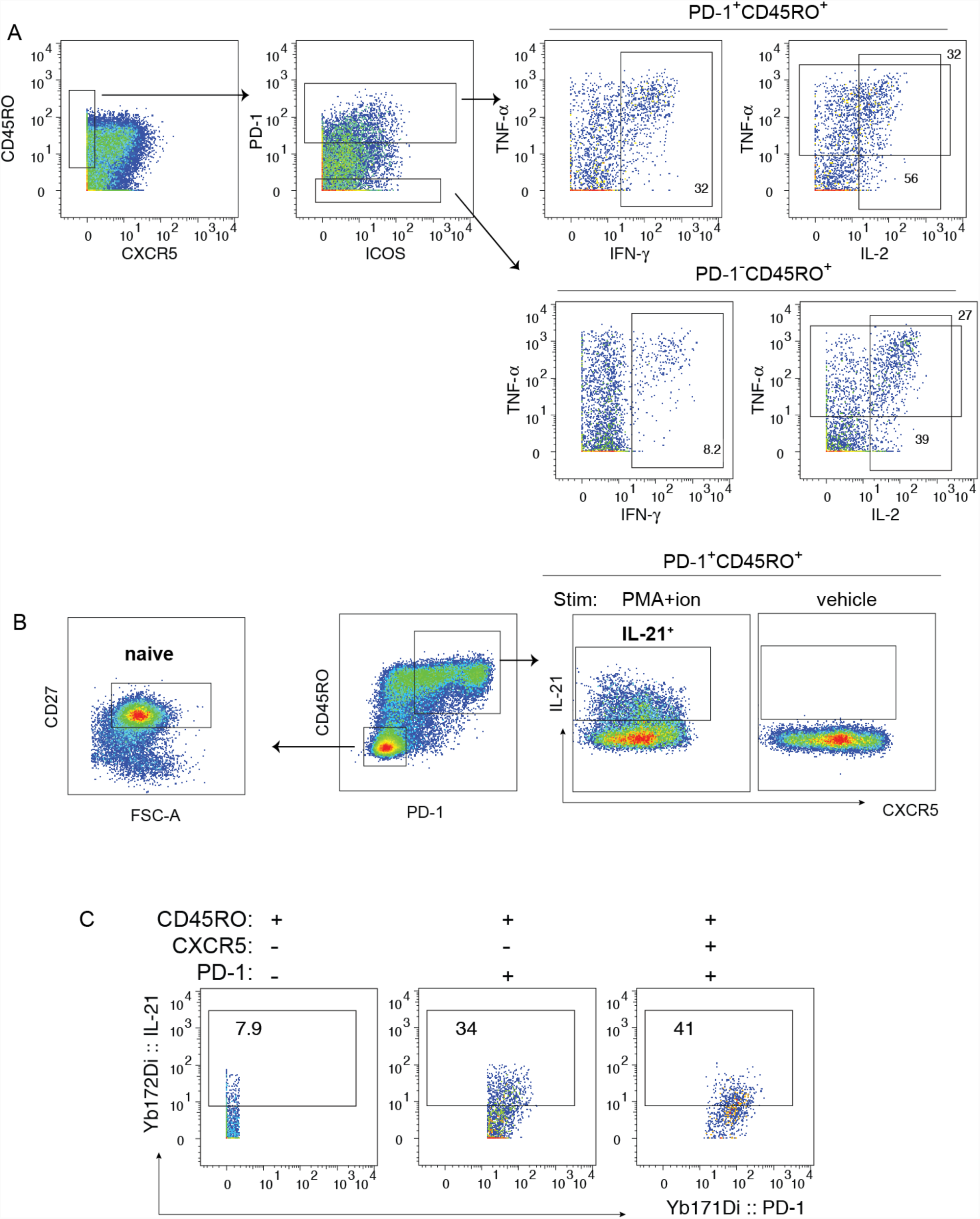
Identification of cytokine producing CD4^+^ T cells. (A) Example plots showing Boolean gates applied on PD-1^+^ and PD-1^-^ memory T cells for the indicated cytokines. (B) Representative plots showing IL-21 expression on PD-1^+^ memory T cells with or without stimulation by PMA and ionomycin. Gates also include example plot for naïve cells. (C) Plots showing representative IL-21 staining on each indicated cell subset.

**Figure S5:**
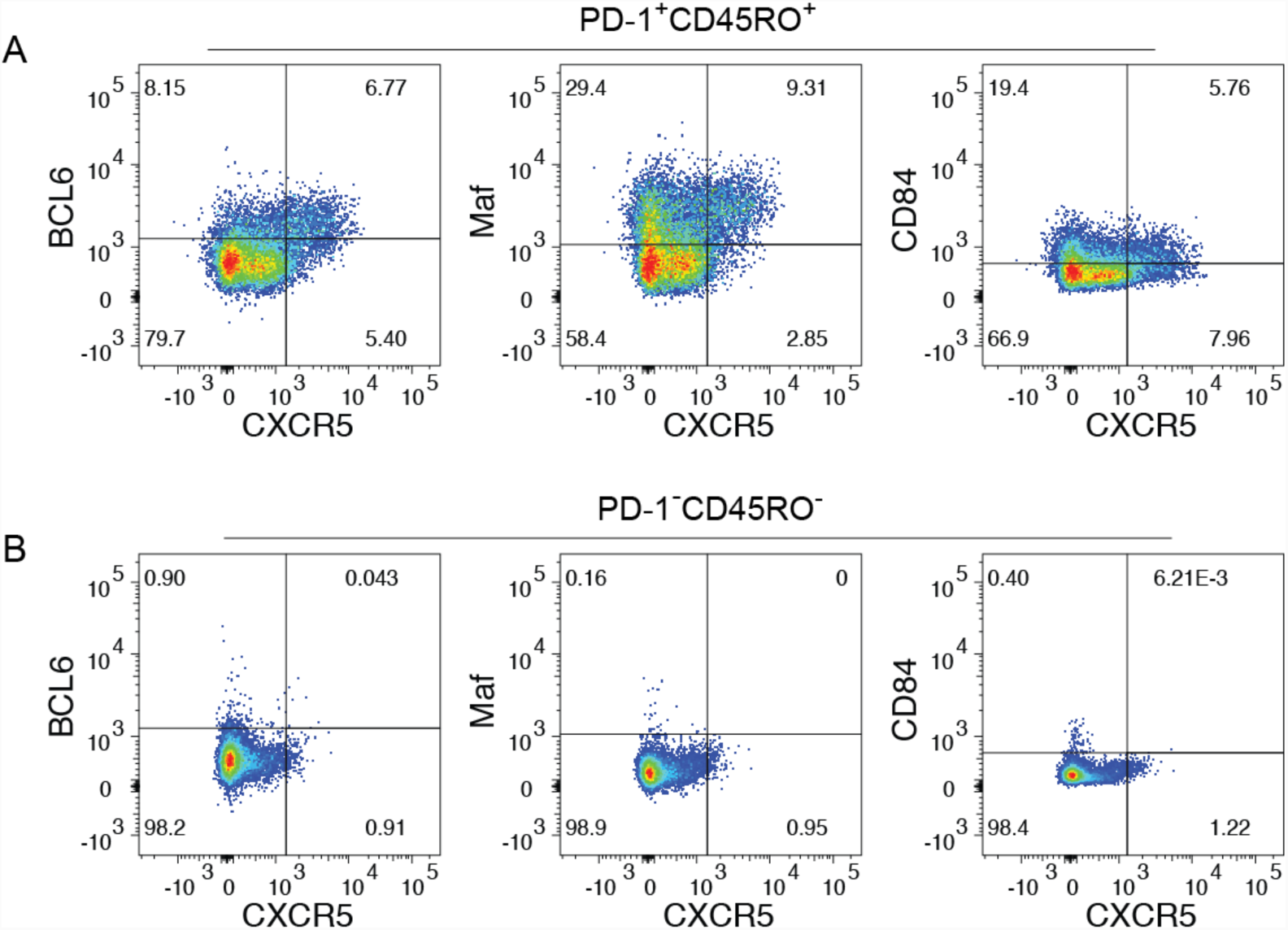
Expression of BCL6, Maf, and CD84 on CD4^+^ T cells. (A-B) Representative plots showing staining for BCL6, Maf, and CD84 on PD-1^+^CD45RO^+^CD4^+^ T cells (A) from HIV infected LNs. Identical gates were applied on PD-1^-^CD45RO^-^ naïve CD4^+^ T cells for comparison (B).

**Figure S6:**
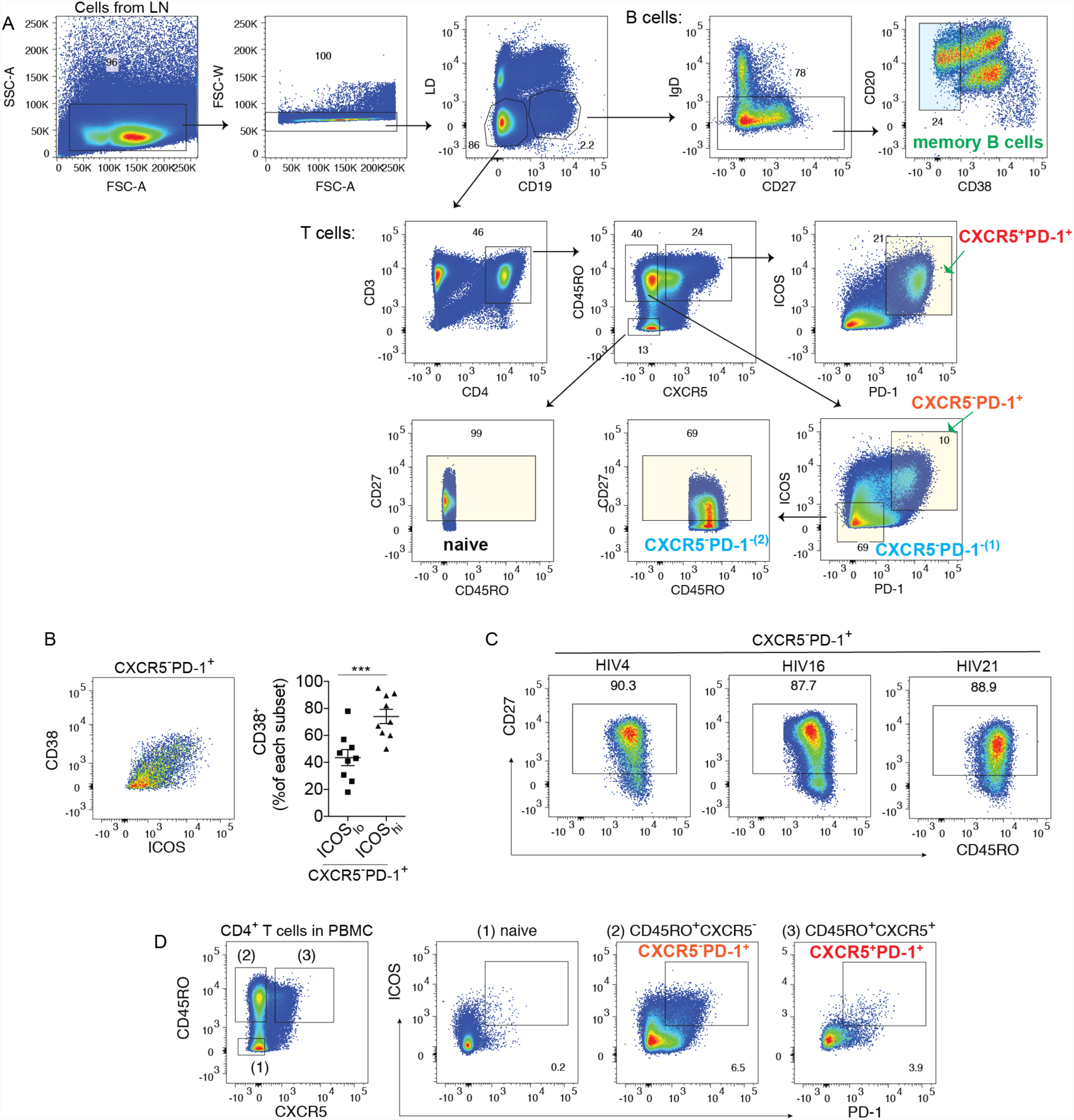
Sort gates for the isolation of T cell and B cell subsets. (A) Representative gates used for sorting memory B cells and T cell subsets. Memory B cells, CXCR5^+^PD-1^+^ T cells, and CXCR5^-^PD-1^+^ T cells from the LNs were sorted for coculture assays. For sequencing experiments (TCR, RNA, ATAC-seq), naïve and CXCR5^-^PD-1^-^ were also sorted as indicated. CXCR5^-^PD-1^-^ T cells sorted for TCR sequencing were gated on ICOS^-^PD-1^-^ subset (CXCR5^-^PD-1^- (1)^). CXCR5^-^PD-1^-^ T cells were additionally gated on CD27 to define central memory cells for RNA and ATAC-seq experiments (CXCR5^-^PD-1^- (2)^). (B) Representative plot showing CD38 and ICOS staining on CXCR5^-^PD-1^+^ T cells. Scatter plot quantifies CD38^+^ T cells and shows enrichment for activated CXCR5^-^PD-1^+^ T cells in cells expressing high levels of ICOS. (C) CD27 antibody stained the majority of CXCR5^-^PD-1^+^ T cells. Plots show CD27 expression on CXCR5^-^PD-1^+^ T cells from samples used for transcriptomic and epigenetic analyses. (D) ICOS^+^PD-1^+^ phenotype was less distinct in CXCR5^+^PD-1^+^ T cells in the PBMC and therefore positive ICOS^+^PD-1^+^ gates were set by the absence of these cells in the naïve population. Example plot of ICOS^+^PD-1^+^ gate used for sorting CXCR5^-^PD-1^+^ and CXCR5^+^PD-1^+^ T cells in the PBMCs. For (B), paired t-test was performed.

## References

Arnold, C.N., Campbell, D.J., Lipp, M., and Butcher, E.C. (2007). The germinal center response is impaired in the absence of T cell-expressed CXCR5. Eur J Immunol 37, 100–109.

Balagopal, A., Asmuth, D.M., Yang, W.T., Campbell, T.B., Gupte, N., Smeaton, L., Kanyama, C., Grinsztejn, B., Santos, B., Supparatpinyo, K., et al. (2015). Pre-cART Elevation of CRP and CD4+ T-Cell Immune Activation Associated With HIV Clinical Progression in a Multinational Case-Cohort Study. J Acquir Immune Defic Syndr 70, 163–171.

Bauquet, A.T., Jin, H., Paterson, A.M., Mitsdoerffer, M., Ho, I.C., Sharpe, A.H., and Kuchroo, V.K. (2009). The costimulatory molecule ICOS regulates the expression of c-Maf and IL-21 in the development of follicular T helper cells and TH-17 cells. Nat Immunol 10, 167–175.

Bentebibel, S.E., Lopez, S., Obermoser, G., Schmitt, N., Mueller, C., Harrod, C., Flano, E., Mejias, A., Albrecht, R.A., Blankenship, D., et al. (2013). Induction of ICOS+CXCR3+CXCR5+ TH cells correlates with antibody responses to influenza vaccination. Science translational medicine 5, 176ra132.

Bhattacharyya, A. (1943). On a measure of divergence between two statistical populations defined by their probability distribution. Bulletin of the Calcutta Mathematical Society 35, 99–110.

Biancotto, A., Grivel, J.C., Iglehart, S.J., Vanpouille, C., Lisco, A., Sieg, S.F., Debernardo, R., Garate, K., Rodriguez, B., Margolis, L.B., and Lederman, M.M. (2007). Abnormal activation and cytokine spectra in lymph nodes of people chronically infected with HIV-1. Blood 109, 4272–4279.

Cannons, J.L., Qi, H., Lu, K.T., Dutta, M., Gomez-Rodriguez, J., Cheng, J., Wakeland, E.K., Germain, R.N., and Schwartzberg, P.L. (2010). Optimal germinal center responses require a multistage T cell:B cell adhesion process involving integrins, SLAM-associated protein, and CD84. Immunity 32, 253–265.

Chevalier, N., Jarrossay, D., Ho, E., Avery, D.T., Ma, C.S., Yu, D., Sallusto, F., Tangye, S.G., and Mackay, C.R. (2011). CXCR5 expressing human central memory CD4 T cells and their relevance for humoral immune responses. J Immunol 186, 5556–5568.

Crawford, A., Angelosanto, J.M., Kao, C., Doering, T.A., Odorizzi, P.M., Barnett, B.E., and Wherry, E.J. (2014). Molecular and Transcriptional Basis of CD4(+) T Cell Dysfunction during Chronic Infection. Immunity.

Crotty, S. (2014). T follicular helper cell differentiation, function, and roles in disease. Immunity 41, 529–542.

Crum-Cianflone, N.F., Iverson, E., Defang, G., Blair, P.J., Eberly, L.E., Maguire, J., Ganesan, A., Faix, D., Duplessis, C., Lalani, T., et al. (2011). Durability of antibody responses after receipt of the monovalent 2009 pandemic influenza A (H1N1) vaccine among HIV-infected and HIV-uninfected adults. Vaccine 29, 3183–3191.

Cubas, R.A., Mudd, J.C., Savoye, A.L., Perreau, M., van Grevenynghe, J., Metcalf, T., Connick, E., Meditz, A., Freeman, G.J., Abesada-Terk, G., Jr., et al. (2013). Inadequate T follicular cell help impairs B cell immunity during HIV infection. Nat Med 19, 494–499.

Cyster, J.G., and Schwab, S.R. (2012). Sphingosine-1-phosphate and lymphocyte egress from lymphoid organs. Annu Rev Immunol 30, 69–94.

Dan, J.M., Lindestam Arlehamn, C.S., Weiskopf, D., da Silva Antunes, R., Havenar-Daughton, C., Reiss, S.M., Brigger, M., Bothwell, M., Sette, A., and Crotty, S. (2016). A Cytokine-Independent Approach To Identify Antigen-Specific Human Germinal Center T Follicular Helper Cells and Rare Antigen-Specific CD4+ T Cells in Blood. J Immunol 197, 983–993.

de Armas, L.R., Pallikkuth, S., George, V., Rinaldi, S., Pahwa, R., Arheart, K.L., and Pahwa, S. (2017). Reevaluation of immune activation in the era of cART and an aging HIV-infected population. JCI insight 2.

Djuretic, I.M., Levanon, D., Negreanu, V., Groner, Y., Rao, A., and Ansel, K.M. (2007). Transcription factors T-bet and Runx3 cooperate to activate Ifng and silence Il4 in T helper type 1 cells. Nat Immunol 8, 145–153.

Finck, R., Simonds, E.F., Jager, A., Krishnaswamy, S., Sachs, K., Fantl, W., Pe’er, D., Nolan, G.P., and Bendall, S.C. (2013). Normalization of mass cytometry data with bead standards. Cytometry A 83, 483–494.

Giorgi, J.V., Hultin, L.E., McKeating, J.A., Johnson, T.D., Owens, B., Jacobson, L.P., Shih, R., Lewis, J., Wiley, D.J., Phair, J.P., et al. (1999). Shorter survival in advanced human immunodeficiency virus type 1 infection is more closely associated with T lymphocyte activation than with plasma virus burden or virus chemokine coreceptor usage. J Infect Dis 179, 859–870.

Giorgi, J.V., Liu, Z., Hultin, L.E., Cumberland, W.G., Hennessey, K., and Detels, R. (1993). Elevated levels of CD38+ CD8+ T cells in HIV infection add to the prognostic value of low CD4+ T cell levels: results of 6 years of follow-up. The Los Angeles Center, Multicenter AIDS Cohort Study. J Acquir Immune Defic Syndr 6, 904–912.

Haghverdi, L., Buettner, F., and Theis, F.J. (2015). Diffusion maps for high-dimensional single-cell analysis of differentiation data. Bioinformatics 31, 2989–2998.

Hale, J.S., Youngblood, B., Latner, D.R., Mohammed, A.U., Ye, L., Akondy, R.S., Wu, T., Iyer, S.S., and Ahmed, R. (2013). Distinct memory CD4+ T cells with commitment to T follicular helper- and T helper 1-cell lineages are generated after acute viral infection. Immunity 38, 805–817.

Hao, Y., O’Neill, P., Naradikian, M.S., Scholz, J.L., and Cancro, M.P. (2011). A B-cell subset uniquely responsive to innate stimuli accumulates in aged mice. Blood 118, 1294–1304.

Havenar-Daughton, C., Lee, J.H., and Crotty, S. (2017). Tfh cells and HIV bnAbs, an immunodominance model of the HIV neutralizing antibody generation problem. Immunol Rev 275, 49–61.

Haynes, N.M., Allen, C.D., Lesley, R., Ansel, K.M., Killeen, N., and Cyster, J.G. (2007). Role of CXCR5 and CCR7 in follicular Th cell positioning and appearance of a programmed cell death gene-1high germinal center-associated subpopulation. J Immunol 179, 5099–5108.

He, J., Tsai, L.M., Leong, Y.A., Hu, X., Ma, C.S., Chevalier, N., Sun, X., Vandenberg, K., Rockman, S., Ding, Y., et al. (2013). Circulating precursor CCR7(lo)PD-1(hi) CXCR5(+) CD4(+) T cells indicate Tfh cell activity and promote antibody responses upon antigen reexposure. Immunity 39, 770–781.

Hong, J.J., Chang, K.T., and Villinger, F. (2016). The Dynamics of T and B Cells in Lymph Node during Chronic HIV Infection: TFH and HIV, Unhappy Dance Partners? Frontiers in immunology 7, 522.

Hufert, F.T., van Lunzen, J., Janossy, G., Bertram, S., Schmitz, J., Haller, O., Racz, P., and von Laer, D. (1997). Germinal centre CD4+ T cells are an important site of HIV replication in vivo. AIDS 11, 849–857.

Hunt, P.W., Lee, S.A., and Siedner, M.J. (2016). Immunologic Biomarkers, Morbidity, and Mortality in Treated HIV Infection. J Infect Dis 214 Suppl 2, S44–50.

Karim, R., Mack, W.J., Stiller, T., Operskalski, E., Frederick, T., Landay, A., Young, M.A., Tien, P.C., Augenbraun, M., Strickler, H.D., and Kovacs, A. (2013). Association of HIV clinical disease progression with profiles of early immune activation: results from a cluster analysis approach. AIDS 27, 1473–1481.

Ke-Yue Ma, C.H., Ben S. Wendel, Chad M. Williams, Jun Xiao, Hui Yang, and Ning Jiang (2018). Immune Repertoire Sequencing using Molecular Identifiers Enables Accurate Clonality Discovery and Clone Size Quantification. Frontiers in immunology In press.

Knox, J.J., Buggert, M., Kardava, L., Seaton, K.E., Eller, M.A., Canaday, D.H., Robb, M.L., Ostrowski, M.A., Deeks, S.G., Slifka, M.K., et al. (2017). T-bet+ B cells are induced by human viral infections and dominate the HIV gp140 response. JCI insight 2.

Kohler, S.L., Pham, M.N., Folkvord, J.M., Arends, T., Miller, S.M., Miles, B., Meditz, A.L., McCarter, M., Levy, D.N., and Connick, E. (2016). Germinal Center T Follicular Helper Cells Are Highly Permissive to HIV-1 and Alter Their Phenotype during Virus Replication. J Immunol 196, 2711–2722.

Langford, S.E., Ananworanich, J., and Cooper, D.A. (2007). Predictors of disease progression in HIV infection: a review. AIDS research and therapy 4, 11.

Levine, J.H., Simonds, E.F., Bendall, S.C., Davis, K.L., Amirel, A.D., Tadmor, M.D., Litvin, O., Fienberg, H.G., Jager, A., Zunder, E.R., et al. (2015). Data-Driven Phenotypic Dissection of AML Reveals Progenitor-like Cells that Correlate with Prognosis. Cell 162, 184–197.

Lindqvist, M., van Lunzen, J., Soghoian, D.Z., Kuhl, B.D., Ranasinghe, S., Kranias, G., Flanders, M.D., Cutler, S., Yudanin, N., Muller, M.I., et al. (2012). Expansion of HIV-specific T follicular helper cells in chronic HIV infection. J Clin Invest 122, 3271–3280.

Locci, M., Havenar-Daughton, C., Landais, E., Wu, J., Kroenke, M.A., Arlehamn, C.L., Su, L.F., Cubas, R., Davis, M.M., Sette, A., et al. (2013). Human circulating PD-1+CXCR3-CXCR5+ memory Tfh cells are highly functional and correlate with broadly neutralizing HIV antibody responses. Immunity 39, 758–769.

Lorenzo-Redondo, R., Fryer, H.R., Bedford, T., Kim, E.Y., Archer, J., Kosakovsky Pond, S.L., Chung, Y.S., Penugonda, S., Chipman, J.G., Fletcher, C.V., et al. (2016). Persistent HIV-1 replication maintains the tissue reservoir during therapy. Nature 530, 51–56.

Matthieu, P., Anne-Laure, S., Elisa De, C., Jean-Marc, C., Rafael, C., Elias, K.H., Laurence De, L., Cecilia, G., and Giuseppe, P. (2013). Follicular helper T cells serve as the major CD4 T cell compartment for HIV-1 infection, replication, and production. J Exp Med 210, 143–156.

Moir, S., Ho, J., Malaspina, A., Wang, W., DiPoto, A.C., O’Shea, M.A., Roby, G., Kottilil, S., Arthos, J., Proschan, M.A., et al. (2008). Evidence for HIV-associated B cell exhaustion in a dysfunctional memory B cell compartment in HIV-infected viremic individuals. J Exp Med 205, 1797–1805.

Morita, R., Schmitt, N., Bentebibel, S.E., Ranganathan, R., Bourdery, L., Zurawski, G., Foucat, E., Dullaers, M., Oh, S., Sabzghabaei, N., et al. (2011). Human blood CXCR5(+)CD4(+) T cells are counterparts of T follicular cells and contain specific subsets that differentially support antibody secretion. Immunity 34, 108–121.

Naradikian, M.S., Myles, A., Beiting, D.P., Roberts, K.J., Dawson, L., Herati, R.S., Bengsch, B., Linderman, S.L., Stelekati, E., Spolski, R., et al. (2016). Cutting Edge: IL-4, IL-21, and IFN-gamma Interact To Govern T-bet and CD11c Expression in TLR-Activated B Cells. J Immunol 197, 1023–1028.

Nish, S.A., Zens, K.D., Kratchmarov, R., Lin, W.W., Adams, W.C., Chen, Y.H., Yen, B., Rothman, N.J., Bhandoola, A., Xue, H.H., et al. (2017). CD4+ T cell effector commitment coupled to self-renewal by asymmetric cell divisions. J Exp Med 214, 39–47.

Rao, D.A., Gurish, M.F., Marshall, J.L., Slowikowski, K., Fonseka, C.Y., Liu, Y., Donlin, L.T., Henderson, L.A., Wei, K., Mizoguchi, F., et al. (2017). Pathologically expanded peripheral T helper cell subset drives B cells in rheumatoid arthritis. Nature 542, 110–114.

Rimpei, M., Nathalie, S., Salah-Eddine, B., Rajaram, R., Laure, B., Gerard, Z., Emile, F., Melissa, D., SangKon, O., Natalie, S., et al. (2011). Human blood CXCR5(+)CD4(+) T cells are counterparts of T follicular cells and contain specific subsets that differentially support antibody secretion. Immunity 34, 108–121.

Rubtsova, K., Rubtsov, A.V., van Dyk, L.F., Kappler, J.W., and Marrack, P. (2013). T-box transcription factor T-bet, a key player in a unique type of B-cell activation essential for effective viral clearance. Proc Natl Acad Sci U S A 110, E3216–3224.

Sereti, I., and Altfeld, M. (2016). Immune activation and HIV: an enduring relationship. Curr Opin HIV AIDS 11, 129–130.

Shugay, M., Britanova, O.V., Merzlyak, E.M., Turchaninova, M.A., Mamedov, I.Z., Tuganbaev, T.R., Bolotin, D.A., Staroverov, D.B., Putintseva, E.V., Plevova, K., et al. (2014). Towards error-free profiling of immune repertoires. Nature methods.

Simoni, Y., Becht, E., Fehlings, M., Loh, C.Y., Koo, S.L., Teng, K.W.W., Yeong, J.P.S., Nahar, R., Zhang, T., Kared, H., et al. (2018). Bystander CD8(+) T cells are abundant and phenotypically distinct in human tumour infiltrates. Nature 557, 575–579.

Szabo, S.J., Kim, S.T., Costa, G.L., Zhang, X., Fathman, C.G., and Glimcher, L.H. (2000). A novel transcription factor, T-bet, directs Th1 lineage commitment. Cell 100, 655–669.

Wendel, B.S., Del Alcazar, D., He, C., Del Rio-Estrada, P.M., Aiamkitsumrit, B., Ablanedo-Terrazas, Y., Hernandez, S.M., Ma, K.Y., Betts, M.R., Pulido, L., et al. (2018). The receptor repertoire and functional profile of follicular T cells in HIV-infected lymph nodes. Sci Immunol 3.

Yu, D., Rao, S., Tsai, L.M., Lee, S.K., He, Y., Sutcliffe, E.L., Srivastava, M., Linterman, M., Zheng, L., Simpson, N., et al. (2009). The transcriptional repressor Bcl-6 directs T follicular helper cell lineage commitment. Immunity 31, 457–468.

